# Spiders as natural DNA samplers recover arthropod community diversity and biotic interactions across space and time

**DOI:** 10.64898/2026.10.05.756666

**Authors:** Sven Weber, Anja C. Melcher, Jerilyn Calaor, Susan R. Kennedy, Domagoj Gajski, Tim Hörrmann, Arndt Schmidt, Lilly Melzl, Ferdinand Meiß, Julian Hans, Lisa Mahla, Manuel Stothut, Danilo Harms, Klaus Birkhofer, Evan Economo, Haldre Rogers, Rosemary Gillespie, George Roderick, Henrik Krehenwinkel

**Author notes:** Sven Weber and Anja C. Melcher should be considered joint first author.

## Abstract

Global arthropod decline demands effective biodiversity monitoring strategies. However, most current monitoring approaches do not provide an exhaustive picture of arthropod community structure. In particular, biotic interactions and temporal patterns of biodiversity change are still poorly understood. Here we explore the possibility of addressing these two shortfalls using spiders, one of the most important predators of terrestrial arthropods, as natural samplers for arthropod community DNA. We conducted several experiments comparing the recovered community composition between spider gut contents and traditional monitoring methods. Additionally, we used archived spiders from long-term collections to assess the preservation of prey DNA over time. Spiders proved to be highly efficient natural DNA samplers, with gut content metabarcoding revealing similar community composition and α- and β-diversity like traditional methods. Unique arthropod taxa were detected by spider gut contents and traditional methods respectively, indicating that these two approaches can serve as valuable complements to each other. Well-preserved archived spiders effectively reconstructed historical diets up to 20 years back in time, showing that historical spider collections can act as time capsules of the wider diversity that occurred in the spider’s habitat. Spiders as natural samplers overcome critical shortfalls in biodiversity monitoring and contribute to our future understanding of community assembly across space and time.

## 1 Introduction

In light of reports of global arthropod decline (Seibold et al., 2019; Wagner et al., 2021), biodiversity monitoring has become irreplaceable to understand the consequences of environmental change (Outhwaite et al., 2022). Arthropods are commonly monitored using passive trapping devices, which catch highly diverse communities. The development of DNA metabarcoding has revolutionized our ability to taxonomically characterize these communities (Yu et al., 2012) Community metabarcoding allows for the routine assessment of the taxonomic diversity of large bulk community samples of thousands of specimens (Buchner et al., 2023; Krehenwinkel et al., 2019). However, current monitoring programs still leave several challenges unresolved.

A first challenge is understanding the individual species’ ecology (Caron et al., 2022; Rosado et al., 2016). The commonly used passive traps provide little information on species interactions (Busse et al., 2022; Karlsson et al., 2020), which are important indicators for the resilience of ecosystems to environmental change (Jackson & Xue, 2023; Mori et al., 2013). This critical lack of data on biotic interactions has recently been coined the Eltonian Shortfall (Caron et al., 2022; Hortal et al., 2015; Rosado et al., 2016). The missing element is not specimens, but rather information about who interacts with whom, when, and with what intensity (Hortal et al., 2015). A solution to this problem is offered by gut content metabarcoding, which enables the reconstruction of biotic interactions, for example prey-predator relationships, at unprecedented levels of resolution and throughput (Gajski et al., 2024; Krehenwinkel et al., 2017).

A second challenge concerns the lack of standardized time series data of arthropod community diversity. Such data are urgently needed to understand patterns and drivers of biodiversity change (Toju et al., 2024; Ushio et al., 2018). Global insect decline has painfully highlighted the lack of temporal data on arthropod community dynamics (Johnson et al., 2017; van Klink et al., 2020). Research on community trends under global change thus also suffers from a temporal data shortfall (Amendt, 2021; Conrad et al., 2006; Hallmann et al., 2017; Toju et al., 2024; Ushio et al., 2018). This shortfall concerns not only patterns of community diversity, but — even more — biotic interactions and the temporal assembly of food webs.

A possible solution to both issues is dietary analysis of natural sampler organisms (Junk et al., 2025). Besides revealing predator-prey interactions, gut content metabarcoding of generalist consumers can provide crucial information on the taxonomic diversity present in the consumers’ environment. The term “natural sampler” refers to organisms whose tissues are used for the purpose of biodiversity monitoring. Examples include the reconstruction of mammalian biodiversity from gut content of blood-feeding leeches (Siddall et al., 2019) and mosquitoes (Massey et al., 2022) dung beetles (Sigsgaard et al., 2021) or dung flies (Massey et al., 2022; Rodgers et al., 2017), or the reconstruction of aquatic biodiversity from gut contents of detritus-feeding shrimps or filter-feeding mussels and sponges (Junk et al., 2023; Mariani et al., 2019; Siegenthaler et al., 2019; Weber et al., 2023).

For terrestrial arthropod diversity, spiders offer great promise as natural samplers. They are one of the most important predators of terrestrial arthropods and are distributed in almost all terrestrial ecosystems (Nyffeler & Birkhofer, 2017). The majority of spiders are generalist predators, with a diverse diet that is limited only by their ecological traits and prey availability (Birkhofer et al., 2022; Michalko et al., 2019; Potapov et al., 2022). Recent work has already suggested the use of spider webs as broad-scale samplers for community eDNA (Gregorič et al., 2022). Most terrestrial ecosystems harbor diverse spider assemblages belonging to many different hunting guilds (Cardoso et al., 2011; Uetz et al., 1999). By investigating the diets of species that belong to different guilds or specialize on different prey, one could get a near-complete picture of the entire arthropod community. This suggests that spiders as natural samplers will not only provide information on predator-prey interactions but also recover a significant proportion of the taxonomic diversity of arthropod communities.

Besides their potential utility as natural samplers in ongoing or future monitoring efforts, preserved spider specimens may also provide a source for retrospective analyses of biodiversity change, i.e., to overcome the temporal shortfall. Well-preserved natural sampler organisms are excellently suited to reconstruct patterns of biodiversity change over multiple decades (Junk et al., 2023, 2025; Krehenwinkel et al., 2022; Stothut et al., 2024; Weber et al., 2023). Spider samples are commonly stored in ethanol, which is well-suited for long-term DNA preservation. Short DNA sequences can be routinely amplified from ethanol-preserved spiders collected more than a century ago (Krehenwinkel & Pekar, 2015). This suggests that the DNA molecules of prey organisms consumed by the spiders are also preserved during long-term ethanol storage. Enriching and sequencing gut content DNA from spider collections could thus open up novel possibilities to study the assembly of arthropod food webs retrospectively. And using historical spider specimens as natural DNA samplers may even provide information on temporal changes in their prey communities. Analyzing spider gut content could thus close a critical gap in our understanding of temporal and spatial changes in taxonomic and interaction diversity.

Considering the above, here we test two hypotheses: 1) Due to their nature as ubiquitous generalist predators in terrestrial ecosystems, spiders are well-suited natural samplers of arthropod community DNA. Metabarcoding of spider gut content should thus recover a similar taxonomic composition and patterns of α- and β-diversity to traditionally used monitoring approaches. 2) Prey DNA is preserved in the guts of archived spiders and can be characterized by metabarcoding for more than a decade after sampling.

To test these hypotheses, we conducted a series of comparative analyses. We 1a) characterized the taxonomic composition of arthropod communities recovered by spider gut content analysis versus conventional trapping approaches in a German grassland ecosystem. We then 1b) tested the suitability of spiders as natural samplers to recover patterns of α- and β-diversity in forest arthropod communities from three Pacific archipelagos. Using archived spider samples that were either 2a) uncurated or 2b) curated, and corresponding freshly collected samples, we then tested the utility of historical spider collections of varying preservation quality for gut content analysis and biodiversity monitoring. And last, we 3) tested the long-term utility of archived spiders for gut content metabarcoding, using museum spiders collected over the past 150 years.

## 2 Material and methods

### 2.1 Analyzed samples and experiments

To test our hypotheses, we conducted a series of comparative analyses across multiple datasets (hereafter referred to as “experiments” for clarity), each designed to evaluate different aspects of spiders as natural DNA samplers (see Table 1). Each analysis contrasts spider-derived data with independently collected community data or temporal sample sets, rather than representing controlled experimental manipulations.

**Table 1:**
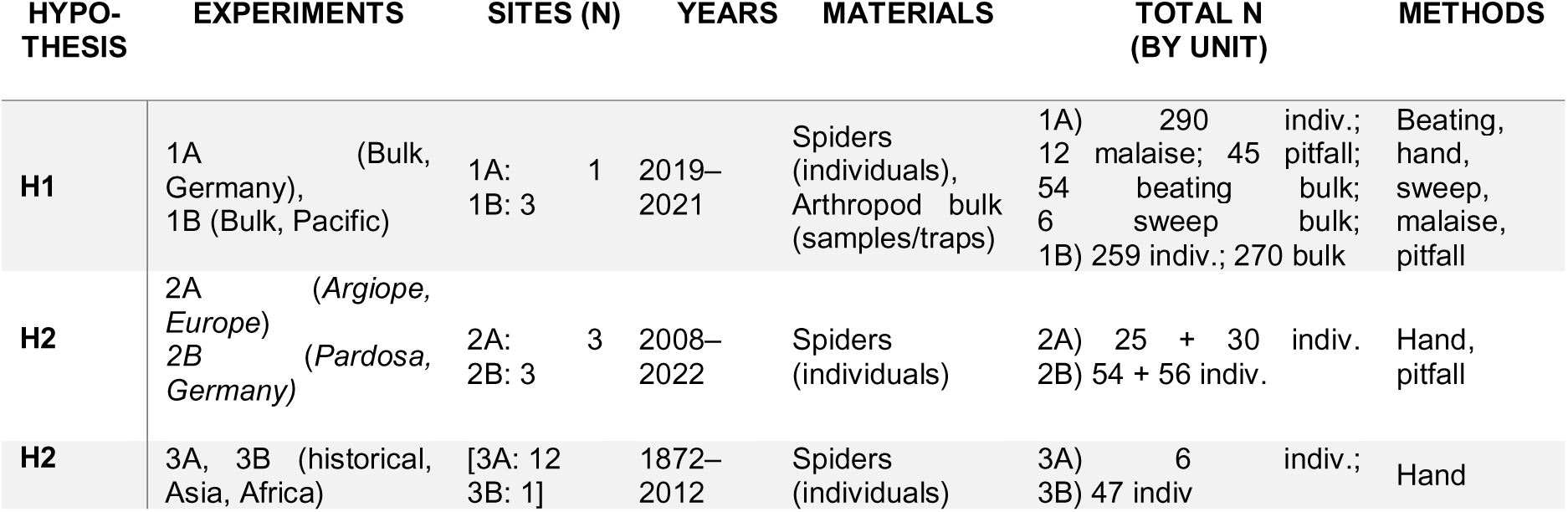
Summary of the spider collections and traditional monitoring regimes. Sample sizes of Arthropod bulk samples refer to: trap samples (containers) for the sampling methods “Malaise Trap” and “Pitfall Trap”, beaten plants for the sample type “Beating” and netting transects for the sample type “Sweep Netting”. Sample sizes for “Spiders” always refer to the numbers of spider individuals, regardless of sampling method. For detailed sampling methods, see (Kennedy et al., 2022; Weber et al., 2024)

| HYPO-THESIS | EXPERIMENTS | SITES (N) | YEARS | MATERIALS | TOTAL N<br>(BY UNIT) | METHODS |
| --- | --- | --- | --- | --- | --- | --- |
| H1 | 1A (Bulk, Germany),<br>1B (Bulk, Pacific) | 1A: 1<br>1B: 3 | 2019–<br>2021 | Spiders (individuals),<br>Arthropod bulk (samples/traps) | 1A) 290 indiv.;<br>12 malaise; 45 pitfall;<br>54 beating bulk;<br>6 sweep bulk;<br>1B) 259 indiv.; 270 bulk | Beating, hand, sweep, malaise, pitfall |
| H2 | 2A (*Argiope*, Europe)<br>2B (*Pardosa*, Germany) | 2A: 3<br>2B: 3 | 2008–<br>2022 | Spiders (individuals) | 2A) 25 + 30 indiv.<br>2B) 54 + 56 indiv. | Hand, pitfall |
| H2 | 3A, 3B (historical, Asia, Africa) | [3A: 12<br>3B: 1] | 1872–<br>2012 | Spiders (individuals) | 3A) 6 indiv.;<br>3B) 47 indiv | Hand |

#### 2.1.1 Experiment 1A: An exhaustive comparison of community diversity recovered by conventional monitoring and spider gut content analysis

For our first comparative analysis, we explored the possibility of obtaining reliable biodiversity data from the gut contents of spiders relative to various traditional arthropod monitoring methods (Table 1). We especially focused on local richness and taxonomic composition recovered by these approaches. Sampling occurred in the summer of 2021 in a nature reserve near Kimmlingen, Germany (49°49’58.4“N 6°36’05.8”E). The site consisted of a semi-dry grassland divided by hedgerows.

Spider specimens were collected by hand, sweep netting in lower vegetation, and beating of hedgerows. Spiders were collected individually and stored in separate tubes filled with pure ethanol. Before processing in the laboratory, individuals were identified to species level under a stereomicroscope. We used 290 specimens of spiders belonging to 19 spider species from 9 families. We chose these specimens because they were collected in high numbers (at least ten specimens per species) with the goal to cover a wide diversity of hunting guilds to maximize the diversity of potential prey. We therefore chose spider species that represented several hunting strategies, such as making specific web types (sheet web, space web, orb web), hunting on the ground or foliage, jumping or ambush hunting (Cardoso et al., 2011). A subset (190 specimens in total) of these 290 specimens has already been published in two studies exploring other methodological aspects, especially pooling and taxonomic saturation (Melcher et al., 2024, 2025). The analyses presented here are distinct and focus on comparing spider-derived community data with conventional sampling methods and evaluating their application within a biomonitoring framework.

For the traditional arthropod monitoring methods, we used samples and data collected during a field study in which traditional sampling was compared to plant derived eDNA (Weber et al., 2024). Briefly, the sampling used a variety of collection methods, including sweep netting, vegetation beating, pitfall and Malaise traps. This sampling is already considerably more exhaustive than most common arthropod monitoring schemes, which usually only rely on one of the above. Arthropods were collected biweekly from three Malaise traps and 15 pitfall traps over six weeks. Vegetation beating for each sample was done by beating three branches of a hedgerow plant, four times each. Sweep netting was executed in transects of 30 m each, by walking back and forth three times with approximately 30 swings per run. These sweeping transects were located left and right of every Malaise trap. Spiders were sampled during the same time period, but independently from bulk samples, with spiders being collected either after or at least 5 days prior to beating or sweep netting of the same plot to avoid cross-contamination and sampling bias due to disturbance. Each spider gut content sample consisted of one spider individual. Adult male spiders were excluded from all experiments because of their reduced feeding activity (Pollard et al., 1995).

#### 2.1.2 Experiment 1B: A comparison of biodiversity metrics recovered by spider gut content and bulk community sampling

In our second experiment, we compared the ability of spider gut content and traditional monitoring to recover patterns of arthropod α-diversity within sites and β-diversity between sites. For this purpose, we used community data from beat sampling in 18 forest sites on islands representing three Pacific archipelagos: the Ryukyus, the Marianas and Hawaii (Kennedy et al., 2022). These samples are part of a large Pacific-wide biomonitoring project in a tropical region and complement the other sampling campaigns in temperate regions. Arthropod samples were collected by beating vegetation at eye level for 20 seconds per sampling point along two parallel transects. One transect was used for bulk community analysis, while spider specimens from the second transect were used for gut content analysis. Spiders from this transect were sorted out, while entire samples from the right transect were used for bulk metabarcoding.

#### 2.1.3 Experiment 2: Exploring the utility of archived spider specimens for gut content analysis

To test whether archived spider specimens still contain amplifiable DNA of their prey, we used two different spider species from different studies: a population genetic study (Krehenwinkel & Tautz, 2013) of the “wasp” spider *Argiope bruennichi,* and a spider community collection of *Pardosa palustris* from the German Biodiversity Exploratories (Fischer et al., 2010), a long-term biodiversity project. Each of these two sample sets consisted of an archived subset of samples (more than ten years old) and a recent subset of samples (sampled in the past 3 years) belonging to the same spider species, collected with the same sampling methods and, in the case of *P. palustris,* from the same sampling sites. The difference between the datasets was the method of preservation: while the *Pardosa* dataset was collected from pitfall traps and the old samples were in poor condition (described below), the *Argiope* specimens were individually hand-collected at both time points and the archived specimens were very well-preserved. Considering their different levels of preservation, we hereafter refer to the two sample sets as uncurated (*Pardosa*) and curated (*Argiope*). Within each set, we distinguish archived (10-year-old samples) and recent (freshly collected samples).

##### Experiment 2A Argiope bruennichi (curated samples)

The first dataset was compiled from specimens of the Palearctic wasp spider *Argiope bruennichi*. The archived dataset comprised 22 specimens from different populations across Europe, which were collected between 2010 and 2012 for a study on the genetics of range expansion (Krehenwinkel & Tautz, 2013). The specimens were individually processed and stored in pure ethanol at room temperature in a room with little to no temperature variation. Ethanol is commonly used for DNA preservation in molecular studies (Taberlet et al., 2018). No visible ethanol evaporation occurred during storage and the specimens’ morphological integrity was very good. These samples served as an example of very good preservation (“curated”). Due to its wide distribution across most of Europe, *A. bruennichi* has access to very different prey communities. To verify the authenticity of the recovered prey DNA, we tested whether the recovered prey species corresponded to species occurring at the respective country of origin. A lack of regional correspondence between a spider’s collection locality and the recovered prey taxa would suggest cross-contamination. As a contemporary comparison, we included 30 specimens of *A. bruennichi* collected in 2022 from Frankfurt (Germany). This dataset allowed us to test the preservation of amplifiable dietary DNA from archived versus freshly collected specimens. To avoid cross-contamination, the DNA extraction and library preparation of historical specimens were performed separately from the recent samples. Using these specimens, we tested whether prey diversity recovered from older preserved spiders is comparable to that found in freshly collected specimens.

##### Experiment 2B Pardosa palustris (uncurated samples)

We analyzed 110 individuals of *Pardosa palustris* from the German Biodiversity Exploratories. We used 54 archived specimens, which were collected during the Soilfoodweb project (Diekötter et al., 2010) with pitfall traps from different field sites in two regions of Germany in 2009 and stored in 70% ethanol at room temperature. As a contemporary comparison, we included 56 specimens from the same locations, also from pitfall traps, but collected more than a decade later in 2021. The archived samples of this comparison have been stored as bulk samples in a university collection since their collection and were not curated according to museum standards. The ethanol had partly evaporated in some samples, with specimens being in poor condition, apparent in extreme cases as some spiders having lost morphological integrity. To avoid cross-contamination, we only included intact spiders in this work (no visible lacerations to prosoma or abdomen, examined under a stereomicroscope) for the analysis. These samples were chosen to explore preservation of prey DNA in badly preserved samples, as is often the case with long-term monitoring samples. The perfect geographic match between these archived and recent samples allowed a direct comparison with reduced influence of geographic differences.

#### 2.1.4 Experiment 3: Exploring the utility of long-term museum collections for gut content metabarcoding

Last, we analyzed spider specimens from a museum collection to further explore the utility of museum spiders for the recovery of prey diversity by gut content metabarcoding. We used spiders of the family Sparassidae (giant huntsman spiders) from the collection of the Senckenberg Museum Frankfurt, Germany. The specimens were collected for taxonomic purposes in tropical or subtropical countries. We analyzed two set of samples from this collection.

##### Experiment 3A Sparassidae contemporary

First, we utilized an unintentional feeding experiment: We included 6 specimens collected between 2004 and 2012. The specimens were all juveniles at the time of collection. Spiders can only be used for taxonomic studies in adult stage, as their genital morphology is not fully developed before this. These specimens were hence reared in the arachnological collection and fed with feeder insects (primarily house crickets) until they reached maturity, and were then preserved for taxonomic study. These specimens thus served as a feeding experiment. We expected that their guts should contain feeder insect DNA, but no wild prey items.

##### Experiment 3B Sparassidae long-term

In addition to the specimens of known diet, we also included additional sparassid specimens from the collection, which were sampled between 1872 and 2012. These specimens served to explore the utility of very old, archived specimens to recover dietary diversity.

### 2.2 DNA isolation

DNA isolation and library preparation of bulk arthropod samples and spiders from Experiment 1, and the archived and freshly collected spiders from experiment 2, was done separately to avoid DNA carryover. Spider specimens for gut content analysis were processed as follows: Before DNA extraction, all spider specimens were individually surface-sterilized with a 0.15% bleach dilution (Greenstone et al., 2012; Miller-ter Kuile et al., 2021). For most DNA extractions, we used whole spider bodies. For *Argiope bruennichi* and the sparassid spiders, which are very large, DNA was extracted only from the opisthosoma. For *Argiope bruennichi*, the whole opisthosoma was used. For Sparassidae, which are larger, we used sterilized forceps to remove tissue from the internal part of the opisthosoma. This was also necessary to preserve the specimens’ integrity for future morphological taxonomic study.

All spider samples were individually homogenized with a bead beater (SPEX 1600MiniG, Metuchen, New Jersey, USA) by adding two sterile 3mm stainless steel beads and lysis buffer followed by shaking at maximum speed for 30 seconds. Afterward, DNA was extracted with the Qiagen Gentra Puregene Cell kit (Qiagen, Hilden, Germany), following the manufacturer’s guidelines. GlycoBlue (Invitrogen, Waltham, United States) was added as coprecipitant during the DNA precipitation step (1:600 volume/volume) to visualize DNA and maximize its yield. DNA extractions were accompanied by blank control extractions to rule out contamination of the reagents or lab environment.

Arthropod bulk samples were sourced from Weber et al. (2024) (experiment 1A) and Kennedy et al. (2022) (experiment 1B). They were extracted using a non-destructive approach, also using the Qiagen Gentra Puregene kit according to the manufacturer’s protocol. Bulk samples were lysed overnight in 200 µl - 250 ml of lysis buffer, depending on biomass. For large animals over 20 mm body size, only a leg was added to the bulk samples before starting the extraction to reduce their contribution to the overall DNA sample. 600 µl of each lysate was then used for DNA extraction (Porco et al., 2010).

### 2.3 PCR amplification, library preparation and sequencing

We used four pairs of primers (Supplementary table S1), one targeting a 115 bp region of the mitochondrial COI gene (Experiments 1A, 2A, 2B) (Spidprey_F/Spidprey_R; Melcher et al. 2) and three (Experiment 1B) targeting fragments of 120-250 bp of the nuclear 18S and 28S rDNA (Krehenwinkel et al., 2019). The latter markers evolve significantly more slowly than COI and hence are not always suited to recover species-level assignments. However, they have been shown to recover very similar taxonomic composition and patterns of biodiversity to standard COI barcoding markers (de Kerdrel et al., 2020; Krehenwinkel et al., 2019). Using three independent primer pairs, each with unique biases in a multiplex, yields a very complete prey community (Krehenwinkel et al., 2019). The fragments amplified by all primer pairs are very short, accounting for prey DNA degradation in the spider’s gut. Such short fragment lengths have been shown to amplify reliably from spiders archived in museum collections for extended periods (Krehenwinkel & Pekar, 2015). All primer pairs were designed to minimize spider DNA amplification while amplifying a broad spectrum of other arthropods. This way, trace amounts of prey DNA can be enriched from DNA extracts that are heavily dominated by spider DNA.

Bulk community samples, as well as spider gut content samples, were amplified with the same primer pairs to make them comparable. PCRs were performed using the Qiagen Multiplex PCR kit with an annealing temperature of 46 °C for the mitochondrial and 55 °C for the nuclear markers and with 32 cycles in 10µl volumes according to the manufacturer’s protocol. For all markers, one blank PCR reaction (with nuclease-free water instead of template DNA) was included as negative control in every PCR batch. All initial PCR primers contained 5’-tails, as described in Lange et al. (2014). On these PCR products, a second 5-cycle dual-indexing PCR was performed to complete a Truseq Library preparation. PCR success was verified on a 1.5% agarose gel. All libraries were then pooled in approximately equimolar amounts based on gel band intensity and the pooled libraries were cleaned of leftover primer using 1X AmPure beads XP (Beckman Coulter, California, USA). The finalized libraries were then sequenced on an Illumina MiSeq using 300-cycle V2 chemistry.

### 2.4 Sequence processing and data analysis

Sequences were demultiplexed using the MiSeq Control Software version 2.6.2.1 with no mismatches allowed. PEAR (Zhang et al., 2014) was used to merge F and R reads, with a minimum quality of 20 and a minimum overlap of 50 bp. FASTX-Toolkit (Gordon & Hannon, 2010) was used to filter all sequences for at least 90% of bases exceeding a quality rating of 30. Sequence processing followed a previously published and validated workflow used in comparable metabarcoding studies, ensuring consistency and direct comparability across datasets (Junk et al., 2023, 2025; Krehenwinkel et al., 2022). While alternative denoising-based pipelines are available, key ecological patterns such as relative diversity and community composition have been shown to be robust across bioinformatic workflows (De Santiago et al., 2022; Glassman & Martiny, 2018). In this study, this is further supported by the consistency of results across clustering levels and taxonomic resolutions (Fig. S3–S5), indicating that the main conclusions are not driven by pipeline-specific artefacts. The quality-filtered sequences were then converted into fasta files and PCR primer sequences were trimmed off from the merged reads using SED and AWK in UNIX. The processed reads were dereplicated and clustered into zero radius operational taxonomic units (zOTUs) using USEARCH including de novo chimera removal (Edgar, 2010). All resulting zOTUs were searched against the NCBI database (downloaded 1/2024) using BLASTn with a maximum of ten target sequences (Altschul et al., 1990). A custom Python script (Schöneberg, 2023) was used to assign taxonomy from the resulting BLAST output. zOTUs with non-arthropod hits among the top ten BLAST hits were excluded from further analyses. We also translated all zOTU sequence and removed those with frameshift mutations or stop codons in the reading frame as likely NUMTS (Graham et al., 2021).. For all others, the top hit was used for taxonomic annotation. Annotated zOTUs were then filtered to a minimum percent identity of 90% and a minimal overlap with the target sequence of 85%. We classified zOTUs to species when they showed a similarity of more than 99% with a reference sequence. A zOTU table was constructed for all samples using USEARCH. Reads below the maximum read number of the respective zOTU in the negative controls were removed. Then, rarefied zOTU tables were created with GUnifrac in R, with the ideal rarefaction depth chosen separately for each dataset, based on the individual saturation and sequencing yield (Fig. S1-2). Rarefied datasets were exported to an Excel worksheet via openxlsx (Schauberger & Walker, 2021). Data were analyzed as relative read abundance per prey taxon to directly compare the results of spider gut content and bulk samples (Deagle et al., 2019).

As zOTUs can inflate recovered taxonomic diversity, we also generated a species-level OTU table, where all zOTUs identified to a single species (≥ 99 % sequence similarity) were merged as one species. This table was used to test for congruence of the diversity patterns recovered by zOTUs with those recovered from actual species. For the slower-evolving nuclear ribosomal markers, we did not identify species, but rather analyzed only at the zOTU level (. However, recent work (de Kerdrel et al., 2020) shows that haplotype-level clustering approximates species status for these markers, making them comparable to our COI dataset in terms of detected diversity patterns. When indicated, we collapsed zOTUs to a single taxon proxy to reduce inflation from micro-variants. We assigned species names when BLAST support met a high-confidence threshold (≥98% identity for species, 100% for higher ranks and sufficient alignment coverage), and otherwise mapped each record to the most specific available rank: genus, family, or order using a simple fallback rule. For readability, unresolved ranks were labeled as “<RANK> spec.” (e.g., Species spec. [*Argiope spec].*, Family spec. [*Lycosidae spec.]*, Order spec. [*Araneae spec.]*). Throughout, we report two richness metrics per sample: (i) zOTU richness and (ii) taxon-collapsed richness (“species richness” sensu this proxy) computed on the collapsed labels to show similarity patterns independent of ASV inflation and to overcome OTU-richness inflation (Fig. S3-5)

### 2.5 Statistical analysis

All data analysis was done in R v 4.2.2 with the package collection of the *tidyverse* (R Core Team, 2021; Wickham et al., 2019). We analyzed the zOTU tables as well as the species-level tables and compared biodiversity patterns between both levels of clustering. Calculation of richness and ordination were performed with *vegan* (Oksanen et al., 2008). Statistical tests were performed with *rstatix* and *report* (Kassambara, 2023; Makowski et al., 2023). Differences in diet composition between spider families and genera of Experiment 1A were each tested via pairwise Permutational Analysis of Variance (PERMANOVA) using the *adonis* function in *vegan*. The difference between genera was additionally tested with pairwise Fisher’s exact tests (Bonferroni corrected). β-diversity of spider gut content samples and bulk samples of Experiment 1B was calculated as pairwise Jaccard dissimilarity between sites. Correlation between spider gut content and bulk samples for richness and β-diversity of experiment 1B was calculated using Pearson correlation.

Differences in amplification success, i.e., number of successfully and unsuccessfully amplified samples of Experiment 2, were tested via chi squared test, followed by a pairwise Fisher’s exact test (Bonferroni corrected) to determine which groups differed. The difference in the proportion of prey DNA versus the spider’s own DNA per sample of experiment 2 was tested using Kruskal-Wallis rank sum tests. Furthermore, Levene’s tests were applied to decipher variation differences between archived and recent samples for both curated and uncurated datasets. To compare the difference in prey composition of archived versus recent *Pardosa* spiders, the relative read abundance of each prey order was compared between the two time points using pairwise Fisher’s exact test with Bonferroni correction.

## 3 RESULTS

### 3.1.1 Experiment 1A: Comparing community diversity recovered by conventional monitoring and spider gut content analysis

To test our first hypothesis that spiders sample a similar arthropod diversity to traditional trapping methods, we compared results from several traditional monitoring methods with spider gut content analysis. The combined spider gut analysis detected a total of 1778 zOTUs (n = 290) and 901 species. Of these, 622 zOTUs and 431 species were unique to spider gut contents, whereas 1156 zOTUs and 527 species were shared with traditional methods. In the pairwise comparison, Malaise trap samples shared 668 of 1620 zOTUs (420 of 1599 species) with spider gut samples (24.5% overlap for zOTUs and 27% overlap for species; Fig. 1A), and pitfall traps shared 270 of 493 zOTUs (166 of 292 species) with spider gut samples (13.5% overlap for zOTUs and 15.2% overlap for species; Fig. 1A,B). This pattern was consistent across clustering approaches and taxonomic resolutions (Fig. S3-S5), and remained similar at higher taxonomic levels (Fig. S6). The relative order composition of the various methods differed slightly: the highest proportion of reads per method belonged to either Diptera, Hemiptera, or Orthoptera, with Malaise trap samples yielding mainly Diptera reads (Fig. 1C). The spider gut content data showed the highest richness in all four individual comparisons and showed higher richness than individual conventional sampling methods (Fig. 1A,B). This held true for both zOTU- and species-level analyses. Traditional methods were sampled to comparable degrees of saturation, whereas spider gut content sampling did not reach full saturation (Fig. S1-S2). About 150 mixed-species spider individuals would need to be analyzed to recover about 2000 zOTUs, whereas approximately 50 beating samples or 10 Malaise traps would be needed to reach the same number (Fig. S2). Netting and pitfall traps did not reach this number of zOTUs in our analysis and were not estimated to do so even with doubled sample size (Fig. S2). Community composition differed significantly among sampling methods (PERMANOVA, Bray–Curtis: F = 9.04, R² = 0.089, p < 0.001). Pairwise comparisons further showed that spider gut content assemblages differed significantly from all conventional sampling approaches (malaise, pitfall, sweep, and beating; all p < 0.001 after correction with BH), confirming consistent compositional differences between spider-derived and trap-based communities. Tests for homogeneity of multivariate dispersion were also significant (betadisper: F = 496.55, p < 0.001), indicating substantial differences in within-method dispersion among sampling approaches. NMDS ordination (stress = 0.184) provided a visualization of these differences, with spider gut content samples showing higher dispersion and overlapping with all traditional sampling methods, whereas conventional methods formed more compact clusters (Fig. S7).

**Figure 1:**
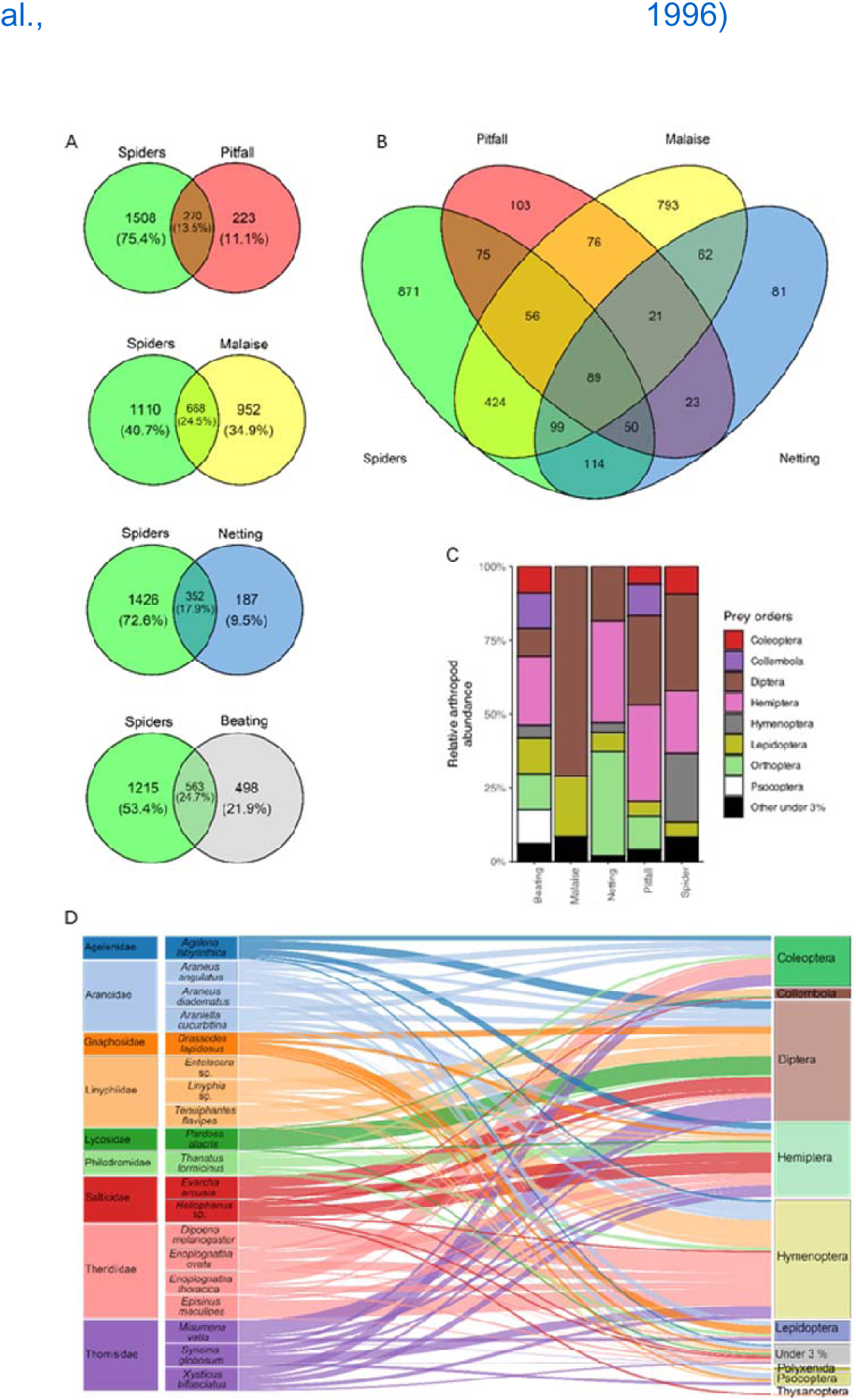
Comparison of taxa recovered from traditional methods versus spider gut content analysis in Experiment 1A. in <u>Germany</u> A) Venn diagrams comparing the number of zOTUs unique and shared between spider gut samples (n = 290) and every traditional method (red = pitfall trap, n = 45; yellow = Malaise trap, n = 12; blue = sweep netting, n = 6; gray = vegetation beating, n = 54). B) Venn diagram comparing the number of zOTUs unique and shared between each tested traditional method and spider gut content analysis (for order level, see Fig. S6). C) Arthropod order composition (Arachnida excluded) detected with each method, based on relative prey read abundance. D) Sankey diagram based on read abundance illustrating the prey-predator network of selected representatives of nine spider families with the ten most abundant prey orders on the right. The thickness of the connections represents the relative number of reads per spider species.

In general, spiders from the grassland in Kimmlingen prey most often on Hymenoptera, Diptera, and Hemiptera. The least commonly detected prey taxa were Polyxenida, Thysanoptera, and Orthoptera, apart from rare taxa amounting to less than 3% read abundance, which were grouped into “other” (Fig. 1D). We found that different spider families have significantly different diets (pairwise PERMANOVA; p < 0.01; Fig. 1D). The analyses even confirmed differences at the genus level (pairwise PERMANOVA; p < 0.01; corrected Fisher’s Exact p adjusted < 0.05). Diptera and Hemiptera prey was consistently abundant in almost all spider families, while other prey orders like Coleoptera, Collembola and Hymenoptera were considerably more likely to be found in certain spider families and far less in others (Fig 1D). While most spider families’ diets contained their main prey orders almost equally abundantly, Theridiidae spiders’ diets were dominated by Hymenoptera prey, with two spider species (*Dipoena melanogaster* and *Episinus maculipes*) almost exclusively feeding on Hymenoptera, as they are described ant specialists (Liu et al., 2022; Umeda et al., 1996).

### 3.1.2 Experiment 1B: A comparison of biodiversity metrics recovered by spider gut content and bulk community sampling

To verify the suitability of spider gut content analysis as a biomonitoring approach to recover large-scale biodiversity patterns, we further tested our protocols in tropical Pacific islands (Experiment 1B). Diversity estimates between bulk arthropod samples collected by beating were highly significantly correlated with those recovered from spider gut content samples. This held true for α-diversity (richness) within sites and β-diversity (Jaccard dissimilarity) between them (R^2^ _richness_= 0.512 (Pearson; F = 22.01; p < 0.01) (Fig. 2A), R^2^_ß-diversity_ = 0.46 (Pearson; F = 54.3; p < 0.01) (Fig. 2B). The order-level composition of spider guts and beating samples was also extremely similar, the main differences being a higher read abundance of Coleoptera and Orthoptera and a lower abundance of Diptera in beating samples than in spider guts (Fig. 2C). This difference, however, was not significant. In detail, bulk arthropod beating samples consisted of 4.6% Coleoptera, 32.0% Collembola, 10.8% Hemiptera, 26.9% Hymenoptera, 6.3% Lepidoptera, 3.4% Orthoptera, 6.7% Psocoptera and 9.2% other orders with under 3% relative read abundance. Spider gut samples consisted of 7.0% Diptera, 36.8% Collembola, 9.3% Hemiptera, 27.5% Hymenoptera, 8.3% Lepidoptera and 11.1% other orders with under 3% abundance (Fig 2C).

**Figure 2:**
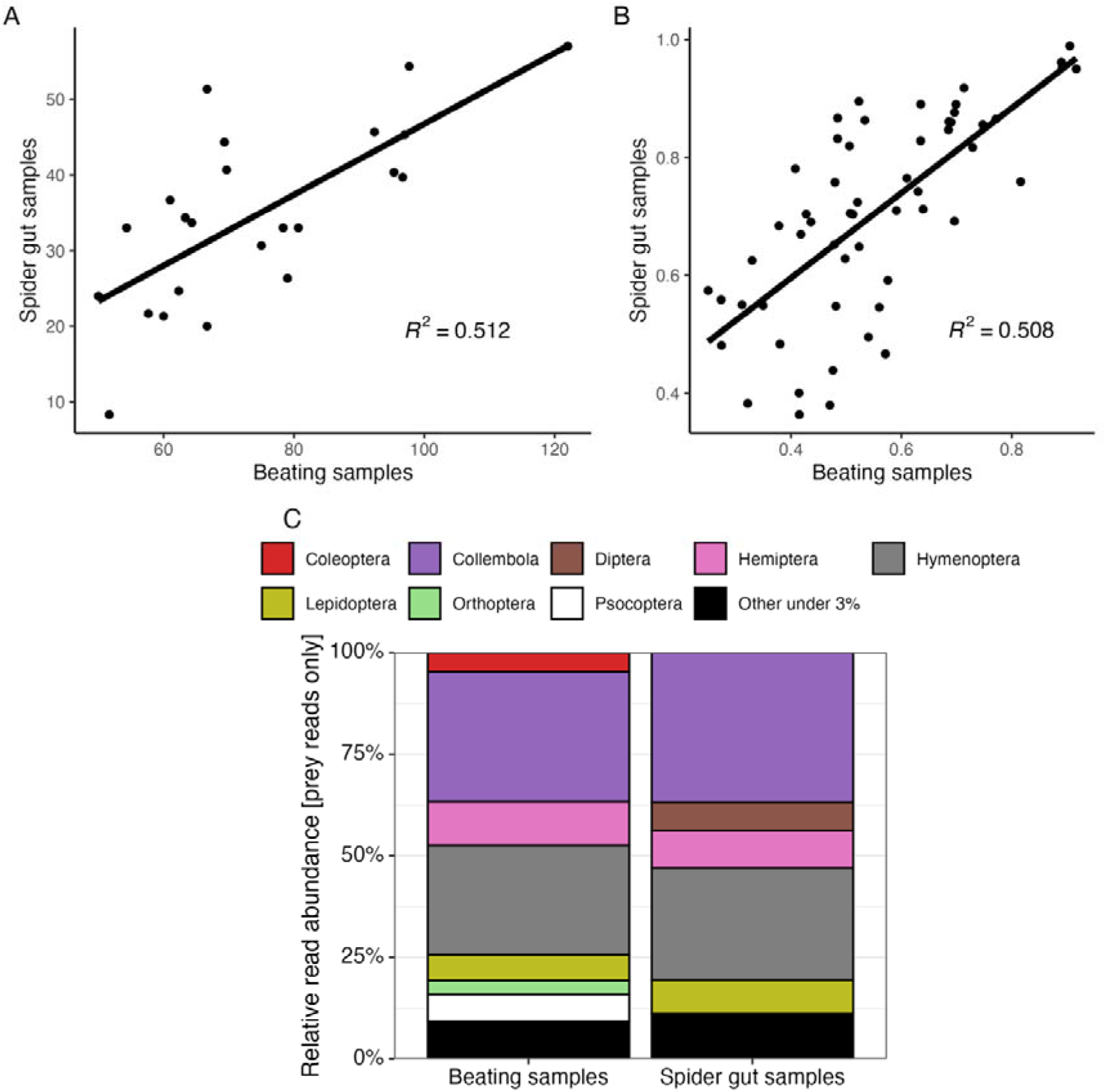
Correlation of diversity estimates for bulk samples from beating versus spider gut content collected from the same transects on <u>three Pacific islands</u>: Okinawa-jima (Ryukyus), Rota (Marianas) and the Big Island (Hawaii). Correlation of (A) zOTU richness and (B) pairwise ß-diversity (Bray-Curtis dissimilarity). C) Stacked barplots showing order-level composition of taxa recovered from bulk beating samples and spider gut samples, based on relative read abundance.

### 3.1.3 Experiment 2: Exploring the utility of archived spider specimens for gut content analysis

We compared the recovery success of prey diversity from archived spiders with that from recently collected ones. This included two datasets representing different levels of long-term sample curation. We first tested the number of samples per dataset for which we successfully recovered prey (> 500 prey reads as sufficient see Fig. S8). For the well-curated spiders (*Argiope bruennichi*), which were preserved in pure ethanol for a decade, we found no significant difference (p > 0.05) in prey recovery success compared to freshly collected specimens of the same species (curated archived: 22/25, curated recent: 30/30). By contrast, a significant difference between archived and freshly collected samples was found for the uncurated samples. The archived *Pardosa palustris* specimens, which were badly preserved, showed prey recovery success for only 23/54 samples, while the freshly collected ones recovered prey for 49/56 (pairwise Fisher’s exact tests, Bonferroni corrected, p <0.001).

We next measured the relative abundance of prey DNA in relation to spider DNA. The relative abundance of prey reads in archived samples was significantly lower than in recently collected samples in both the curated and the uncurated samples (Fig 3B U-test, curated p = 0.001, uncurated p < 0.01). We also found a significantly higher variation of relative prey reads in the archived over the freshly collected samples (Fig 3B Levene’s test, curated p = 0.001, uncurated p < 0.01). This effect was stronger in the uncurated samples where an average of 69% ± 10.97% of prey reads was found for archived specimens and 89.45% ± 30.1% for freshly collected ones. For the curated specimens, we found 99.36% ± 0.97% prey reads for the archived specimens and 99.94% ± 0.09% for the freshly collected ones.

**Figure 3:**
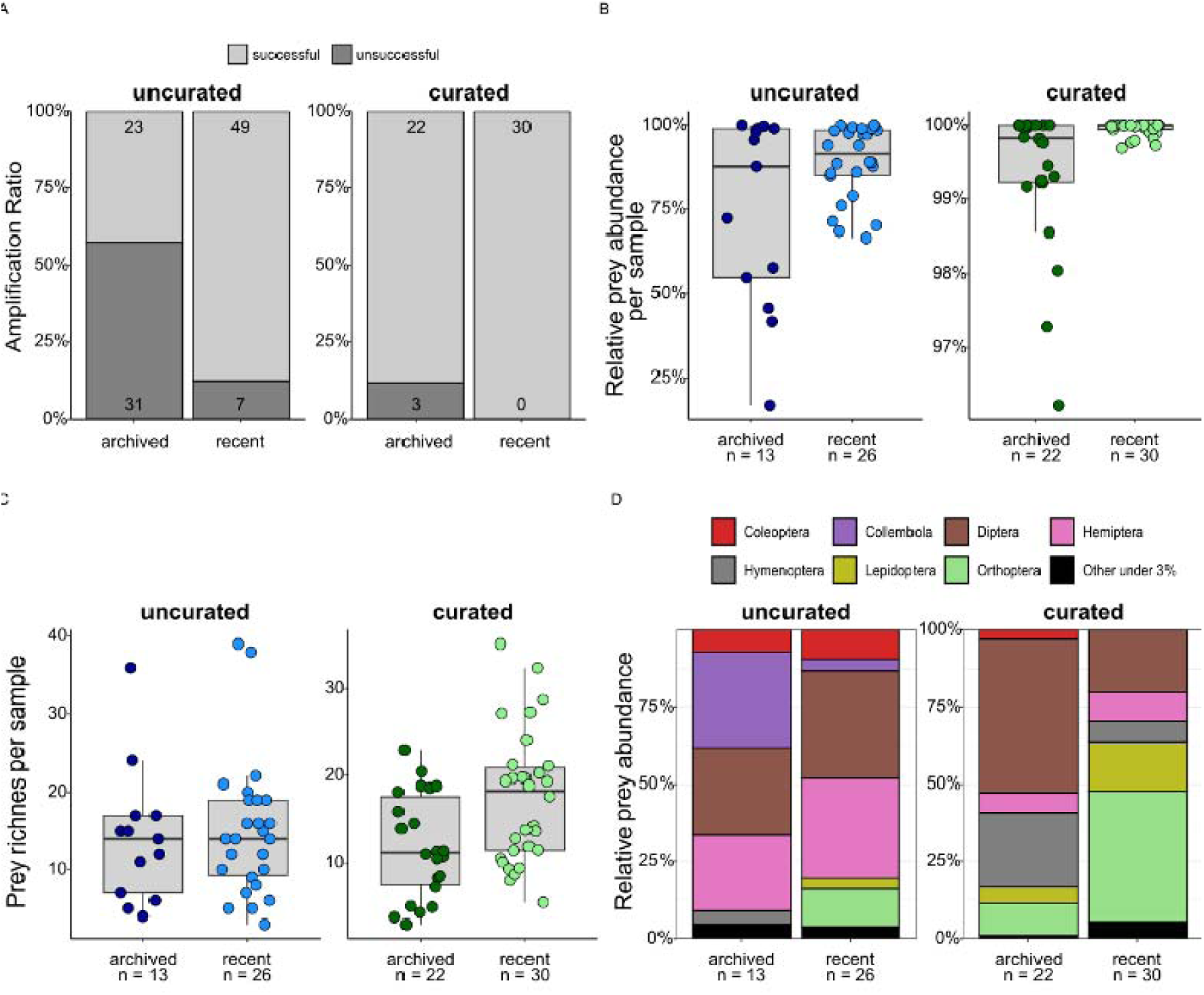
Comparison of the gut contents of archived and freshly collected spider samples (Experiments 2A and 2B) <u>from Germany</u>. (A) Amplification success (calculated as the number of samples exceeding 500 prey reads after quality filtering) for curated archived and recent samples and uncurated recent samples was similar, but significantly lower for uncurated archived samples. (B) Relative read abundance of prey taxa per successful sample (defined as proportion of reads matching arthropod zOTUs that were not conspecific with the predator) was significantly higher in recent samples than in archived staples for both curated and uncurated specimens. Note the different scales on the y-axes for uncurated and curated datasets. (C) Prey (zOTU) richness per sample was slightly lower in both types of archived samples, but not significantly. (D) Relative abundance of the seven most abundant prey orders.

**Figure 4.**
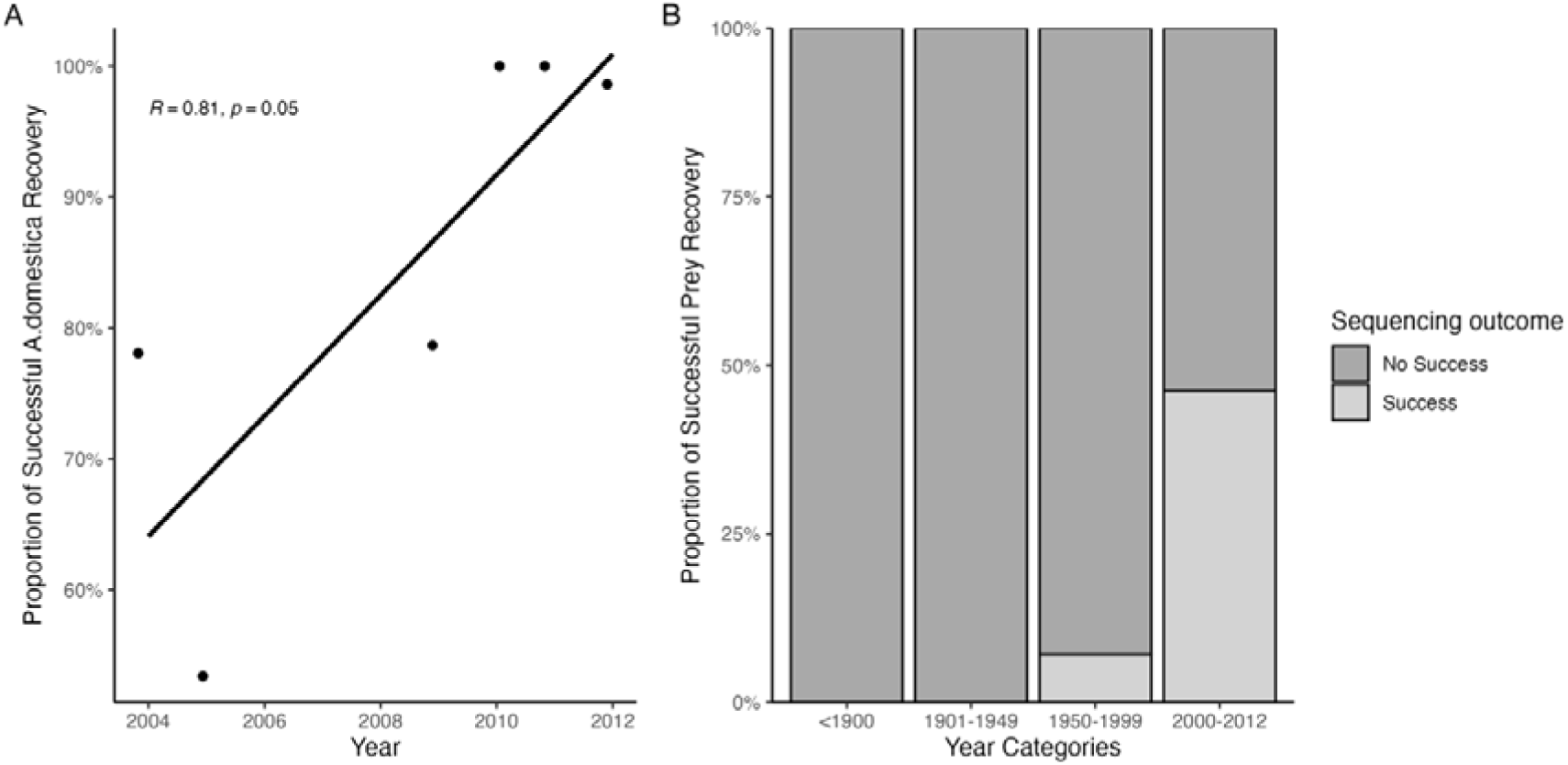
Prey DNA recovery from archived museum spider specimens from Africa and Asia across time. (A) Recovered proportion of expected prey DNA from museum preserved spiders, which were fed in the lab with house crickets, compared against the year in which the spider was added to the collection. The line is a linear regression (Spearman). (B) Amplification success by specimen age class: stacked bars show the proportion of specimens with detectable prey DNA (“Success”) versus no detectable prey DNA (“No Success”).

Next, we analyzed the recovered prey richness for the different datasets. Prey richness did not differ significantly between archived and freshly collected specimens (Fig 3C, Kruskal-Wallis-Test, p > 0.05). This held true for both the curated and uncurated samples and for zOTU richness as well as species richness. We found an average zOTU richness of 11.91 ± 5.97 (species richness = 8.75 ± 4.88) for the curated archived samples and 17.41 ± 7.55 (species richness = 13.82 ± 5.36) for the freshly collected samples of *A. bruennichi*. An even smaller difference was found for the uncurated samples, with an average zOTU richness of 14.96 ± 8.74 (mean species richness = 8.35 ± 5.21) for the archived specimens and 14.07 ± 8.71 (mean species richness = 5.39 ± 2.4) for the freshly collected *P. palustris*.

We also explored dietary composition for the different samples. The analysis of archived and recent samples revealed that *P. palustris* primarily fed on Diptera (31.5% ± 4.95%), Hemiptera (28.56% ± 5.37%), Collembola (17.63% ± 19.5%), Coleoptera (8.16% ± 1.7%), Orthoptera (4.25% ± 8.53%), Hymenoptera (2.06% ± 2.92%) and Lepidoptera (1.73 %± 2.45%). The prey community composition between archived and freshly collected samples was very similar. However, a significant difference was found for Collembola (Fisher’s Exact, p adjusted < 0.01), which showed a higher proportion in the archived spiders, and Orthoptera (Fisher’s Exact, p adjusted < 0.05), which showed a higher proportion in the freshly collected spiders (Fig. 3D).

Archived curated *A. bruennichi* gut content samples, which were sampled from different locations across Europe, mostly contained Diptera (49.55% ± 27.31%), Hymenoptera (24.21% ± 26.26%), Orthoptera (10.28% ± 13.08%), Hemiptera (6.54% ± 7.28%), Lepidoptera (5.23% ± 5.84%) and Coleoptera (3.17% ± 3.68%). Recent curated *A. bruennichi* samples, which, in contrast to the archived ones, were all sampled from the same region in Germany, contained mostly Orthoptera (42.40% ± 35.98%), Diptera (20.10% ± 4.80%), Lepidoptera (15.61% ± 10.90%), Hemiptera (9.51% ± 6.64%) and Hymenoptera (6.90% ± 8.89%). The geographic distribution of all examined zOTUs matched the origin of the respective sample it was detected in (Supplementary table S2).

We then examined whether the detected prey species could be plausibly found in the two spider species. The prey species found in both archived and freshly collected *Pardosa* closely matched the expected prey of this ground hunter (Supplementary table SX). We found typical central European arthropod species, like *Megadelphax sordidulus* (Hemiptera), *Pseudochorthippus parallelus* (Orthoptera) and *Lasius niger* (Hymenoptera). The same held true for the *Argiope bruennichi*: both archived and freshly collected specimens recovered typical prey items for their location of origin (Supplementary table S2). For example, the widespread Central European grasshopper *Chorthippus parallelus* was commonly found in German spiders. In a Portuguese spider, we found the Mediterranean moth species *Luperina dumerilii*, and in an Italian one, the Southern European moth *Scopula minorata*.

### 3.1.4 Experiment 3: Exploring the utility of long-term museum collections for gut content metabarcoding

We last tested six giant huntsman spider specimens from a natural history museum, which were kept in captivity and fed with feeder insects before being preserved between 2004 and 2012. All analyzed specimens yielded detectable prey reads. The used feeder insect was the house cricket *Acheta domestica*, which we found in all specimens in very large read proportions (85 % of the detected prey reads on average). The older the specimens were, the more non-target reads (i.e. reads not belonging to *A. domestica)* were found, likely cross-contamination from the museum collection.

We also tested the utility of even older specimens for gut content analysis, analyzing museum specimens collected between 1872 and 2012. Overall, these specimens worked poorly. None of the specimens collected before 1950 yielded any plausible prey reads. Either no prey reads were recovered, or the recovered reads belonged to prey taxa unlikely to be eaten by the spider. Of 14 specimens collected between 1950 and 1999, only one yielded credible prey information. This was a specimen from Madagascar that seemingly preyed on the Malagasy moth *Panogena lingens*. A considerably larger proportion of specimens yielding interpretable prey information was found for specimens collected after 2000 (46 % of specimens). This included a *Eusparassus* specimen collected in 2004 from the Greek island Lesbos, which likely preyed on *Tettigonia viridissima*, a widespread European bush cricket. A Vietnamese *Pseudopoda* specimen collected in 2005 likely preyed upon the tropical Asian moth *Rhagastis castor.* And a Namibian *Leucorchestris* fed on the moth *Grammodes geometrica,* which is widespread in tropical Africa. Yet, even for specimens collected after 2000, more than 50 % did not yield any prey detections.

## 4 Discussion

### Spiders as natural samplers of arthropod community DNA

Our results confirm our first hypothesis and show that spiders are well suited natural samplers of arthropod community DNA. Spider gut content recovers ecologically meaningful data on community diversity and comparable patterns of α- and β-diversity to those obtained from traditional arthropod sampling methods. But only about half of the recovered arthropod taxa were shared between spider guts and traditional sampling methods. This has important implications for prey–predator studies. Individual trapping approaches are commonly used to identify the available prey for spiders in dietary studies. We show that even multiple traps do not suffice to recover the available prey.

Spiders collected from the field sites may have dispersed from more distant locations, contributing to the wider recovered dietary spectrum. It is also known that prey DNA can be detected back in time for about 2 weeks in a spider (Kennedy et al., 2020). Spider guts may thus cover a time window before the trapping began. The difference in gut content and traditional traps may also be explained by the different DNA isolation protocols we used: destructive for spider guts and non-destructive for trap contents (Iwaszkiewicz-Eggebrecht et al., 2023). However, the non-destructive protocol we used here (Kennedy et al., 2022) has been shown to perform reliably and the combination of several collection methods should decrease possible biases. Moreover, we included arthropod samples from four different trapping approaches, more than 100 trap samples in total. This exhaustive sampling should yield a very accurate picture of the local arthropod diversity. The observed differences between spider gut contents and conventional sampling may also reflect short-term temporal turnover in arthropod communities, as we collected spiders and trap samples on different days.

Overall, our data suggests that spider gut content analysis could serve as a promising complementary approach to traditional sampling. It is well known that different arthropod sampling methods recover different subsets of local diversity (Cuff et al., 2024; Yi et al., 2012; Yu et al., 2012). We do not expect that spider natural samplers will replace traditional arthropod monitoring. But our approach provides an important use for spider bycatch, which can constitute a large proportion of the specimens in bulk traps (Arvidsson et al., 2016; Buchholz et al., 2011; Jennings & Hilburn, 1988; Oxbrough et al., 2010). Often these specimens remain largely unused.

Despite being generalist feeders, spiders can show a certain level of dietary specialization, for example depending on hunting mode. An orb-weaving species with an aerial capture web will recover different prey communities than a ground-hunting species (Birkhofer et al., 2022). To make data from different sites comparable, information about the taxonomy and ecology of the spiders should be incorporated when choosing them as natural samplers. For simple biomonitoring purposes, one could focus on a few widespread generalist spider species (Melcher et al., 2025), which represent diverse life history traits (e.g. different hunting guilds, or body sizes) (Nyffeler & Birkhofer, 2017). Pooling by species can help reduce the necessary workload (Melcher et al. 2024). In some cases, it may even be advantageous to sample specialized spider species. An example could be a study focused on the recovery of ant diversity, sampling specifically the spider species *Dipoena melanogaster* and *Episinus maculipes*, which almost exclusively feed on ants (Fig. 1D).

The utility of spiders as natural sampler has geographical limitations. The spider species composition of different biogeographic regions is very different, with few species being cosmopolitan. Gut content samples from a tropical and temperate ecosystem are thus not be comparable to each other. However, many spiders have a very wide distribution, for example more than 300 spider species are found across the entire Palearctic (Marusik, 1994). Such taxa are well suited to use spiders as natural sampler across entre biogeographic realms.

As spiders are among the most abundant predators of terrestrial invertebrates, they are key elements of terrestrial food webs (Nyffeler & Birkhofer, 2017). A major benefit of spider gut content metabarcoding is that, in addition to revealing arthropod diversity patterns, it provides predator-prey interactions. In contrast, traditional sampling methods only provide taxonomic information on the species present at the site (Busse et al., 2022; Karlsson et al., 2020). Recent literature shows the vast potential of spider gut content DNA for analyzing interaction networks (Casey et al., 2019; Gajski et al., 2024; Suzuki et al., 2023). Using interaction data to construct ecological networks offers the possibility to assess an ecosystem’s robustness and resilience (Pocock et al., 2012).

### The utility of archived spiders for gut content analysis

Our data support our second hypothesis that archived spiders retain their utility for gut content analysis more than a decade after collection. The recovered prey taxa matched the expected prey spectrum of the corresponding spider species very well and reflected the geographic origin of samples. These results support findings by (Cai et al., 2024; Lynggaard et al., 2022; Siddall et al., 2019), which showed the utility of preserved leeches and sponges as natural DNA samplers.

While our results are encouraging, it should be noted that proper storage conditions appear to be very important. We compared two different types of approximately 15-year-old archived spiders: 1) spiders preserved in pure ethanol with no signs of evaporation and 2) spiders stored in 70% ethanol, which had partly evaporated and experienced no curation. Both sample types still provided gut content data, but better-preserved samples did so much more reliably, with nearly all tested specimens still yielding data. This suggests that preservation mode is a critical factor to predict the utility of a specimen. Our data also suggest that the spiders’ own DNA is still amplifiable after the prey DNA is already too degraded for successful amplification (Fig. 3B). Hence, older specimens may show lower measures of prey richness than freshly collected ones, and patterns of prey α-diversity may not be directly comparable. However, the resulting data are still very suitable to study compositional shifts in the spiders’ diets and corresponding arthropod diversity. Provided that the specimens are properly preserved and curated, collecting the same spider species from identical sites over time reveals temporal dynamics of arthropod community composition, trophic specialization and food web structure.

The temporal dynamics of insect populations are still largely understudied (Cardoso et al., 2011; Erwin et al., 2011; Wolfe et al., 1987), primarily due to a lack of standardized temporal data (van Klink et al., 2020, 2022). In addition, most monitoring schemes in the past did not take DNA preservation into account. Often preservatives like formalin were used, leaving samples useless for DNA analysis (Petsopoulos et al., 2024). Here, the use of spiders has another major advantage because they are traditionally stored in ethanol, an excellent DNA preservative. Thus, collections of spiders, especially the numerous unused bycatches from insect monitoring projects, could provide the necessary data to trace changes in insect biodiversity over time, thereby addressing a critical research gap. Further, archived spiders hold great potential to study the temporal dynamics of food webs (Suzuki et al., 2023). Our work on sparassids shows that gut content analysis is also possible by dissecting only a limited amount of tissue from the spider’s digestive tract. This way, the morphological integrity of the specimen can be retained.

Our analysis of museum specimens suggests that PCR-based gut content analysis may not work on specimens considerably older than 20 years. The older the specimen, the less successful amplification and the more potential cross-contamination was recovered. After a certain time in storage, prey DNA may simply be too degraded to be recovered by PCR. Cross-contamination is also very likely in natural history museums, where specimens are often stored in jars with various other species and in ethanol re-used from other specimens. The amount of contaminating DNA sequences may thus increase over time in the museum collection. Future collections should consider storing spiders individually in fresh 95–100% ethanol. This may considerably increase the utility of archived spiders for gut content analysis. The huntsmen spiders we used here, are mostly very large. Previous work suggests that DNA recovery from archived spiders is significantly correlated with the spider’s size (Krehenwinkel & Pekar 2025). This could indicate reduced preservation efficiency in larger individuals, where incomplete submersion or slower ethanol penetration fails to halt digestion quickly.

Our results highlight the need for proper storage, curation and preservation of all collected arthropods from biomonitoring projects, including non-target bycatch like spiders. This could be an important task of natural history collections going forward, and research should be invested into developing suitable curatorial standards to optimize data collection for future biomonitoring collections.

## 5. Conclusions

Our study demonstrates that metabarcoding of spider guts is a powerful tool for assessing arthropod diversity, providing α- and β-diversity patterns comparable to traditional monitoring methods. The marked dietary differences between spider taxa offer several possibilities to utilize spider gut content as a tool for broad diversity surveys or finely resolved arthropod monitoring, optionally combined with interaction networks. We highlight the potential of spider guts as a complementary sampling method, revealing unique prey taxa not captured by traditional techniques. In addition, spider bycatch and archived spider samples are proving valuable for long-term biodiversity studies, provided they are well preserved. These results emphasize the importance of combining spider metabarcoding with conventional methods to improve ecological research, especially to understand trophic interactions and dynamics of arthropod communities across space and time.

## Acknowledgements

This work was partly funded by institutional funds of the Department of Biogeography at Trier University and the DFG Priority Program 1374 “Biodiversity Exploratories” (432985091). We thank Victoria Grießmeier for giving support through the central office and Markus Fischer, Eduard Linsenmair, Dominik Hessenmöller, Daniel Prati, Ingo Schöning, François Buscot, Ernst-Detlef Schulze, Wolfgang W. Weisser and the late Elisabeth Kalko for their role in setting up the Biodiversity Exploratories project. We thank Jonas Rochlitz and Dr. David Ott of the Biodiversity Exploratories project SPRINT for providing the uncurated spider samples of 2020 (Exp. 2A). We thank the Struktur- und Genehmigungsdirektion Nord for granting the necessary permits to conduct sampling at the site in Kimmlingen (425-104-235-005/2021). Sampling in the Pacific was supported by subsidy funding to the Okinawa Institute of Science and Technology and carried out under the approval of the Nature Conservancy and Hawaii Department of Land and Natural Resources (permit no. I1298) and the CNMI Division of Fish and Wildlife. S.W. was supported by the German Federal Environmental Foundation (DBU) [20020/696] and the WB-Fellowship [WE 7675/1-1]. We thank Sven Kuenzel for sequencing our samples. DG was a Brno PhD Talent Scholarship Holder—Funded by the Brno City Municipality. Additionally, for participation in this project, DG was supported by the Operational Programme Research, Development and Education “Project Internal Grant Agency of Masaryk University” (No. CZ.02.2.69 / 0.0 / 0.0 / 19_073 / 0016943). We thank Jordan Cuff and one anonymous reviewer for their helpful feedback on an earlier version of this manuscript.

## Data sharing and availability

All raw reads as well as all zOTU/OTU and metadata tables are available under: https://10.5061/dryad.280gb5n5t.

For the Review please use: https://10.5061/dryad.280gb5n5t-review.

## Declaration of Generative AI and AI-Assisted Technologies in the Writing and Coding Process

During the preparation of this work, the authors used ChatGPT 5 and OpenAI Codex to assist with commenting existing code, refining language, and organizing data and supporting materials for deposition in Dryad. All AI-assisted activities were conducted under human supervision, and the authors reviewed and edited the outputs as needed. The authors take full responsibility for the final content, code accuracy, data organization, and integrity of the published work.

## Author contributions

SW, AM and HK conceived the ideas and designed the study. SW and SK analyzed and visualized the data. SW, AM, JC, SK, DG, TH, AS, LMe, FM, JH, LMa and MS collected and generated the data. DH, KB, EE, HR, SW, RG, GR and HK acquired funding and provided feedback for the final draft of the manuscript. SW led the writing of the manuscript with support of AM.

