## Supplementary material 1 for "Spiders as natural DNA samplers recover arthropod community diversity and biotic interactions across space and time"

**9 SUPPLEMENTARY INFORMATION**

### **Table S1: Primers used for each experiment**. Fragment lengths include primer binding sequences.

| Experiment | Primer | marker type | source | sequence (5’-3’) | fragment length |
| --- | --- | --- | --- | --- | --- |
| 1A, 2AB | Spidprey_F | COI | Melcher et al. (2024) | RGCHTTYCCHCGA HTAAAYAAYA | 115 |
|  | Spidprey_R | COI | Melcher et al. (2024) | RGGRTANACNGTT CAWCCWGT |  |
| 1B | 18S_1392 | 18S | Krehenwinkel et al. 2019 | GTCTGGTTRATTCCG RTAACGAA | 140 |
|  | 18S_1628 | 18S | Krehenwinkel et al. 2019 | CACAGACCTGTTATT GCTCAA |  |
|  | 18S_1332 | 18S | Krehenwinkel et al. 2019 | AGCTCTTTCTYGATT CRGTGGGT | 200 |
|  | 18S_1628 | 18S | Krehenwinkel et al. 2019 | CACAGACCTGTTATT GCTCAA |  |
|  | 28S_788 | 28S | Krehenwinkel et al. 2019 | CGTCTTGAAACACG GACCAA | 250 |
|  | 28S_1062 | 28S | Krehenwinkel et al. 2019 | GWCCTCCATCAGGG TTTCCC |  |

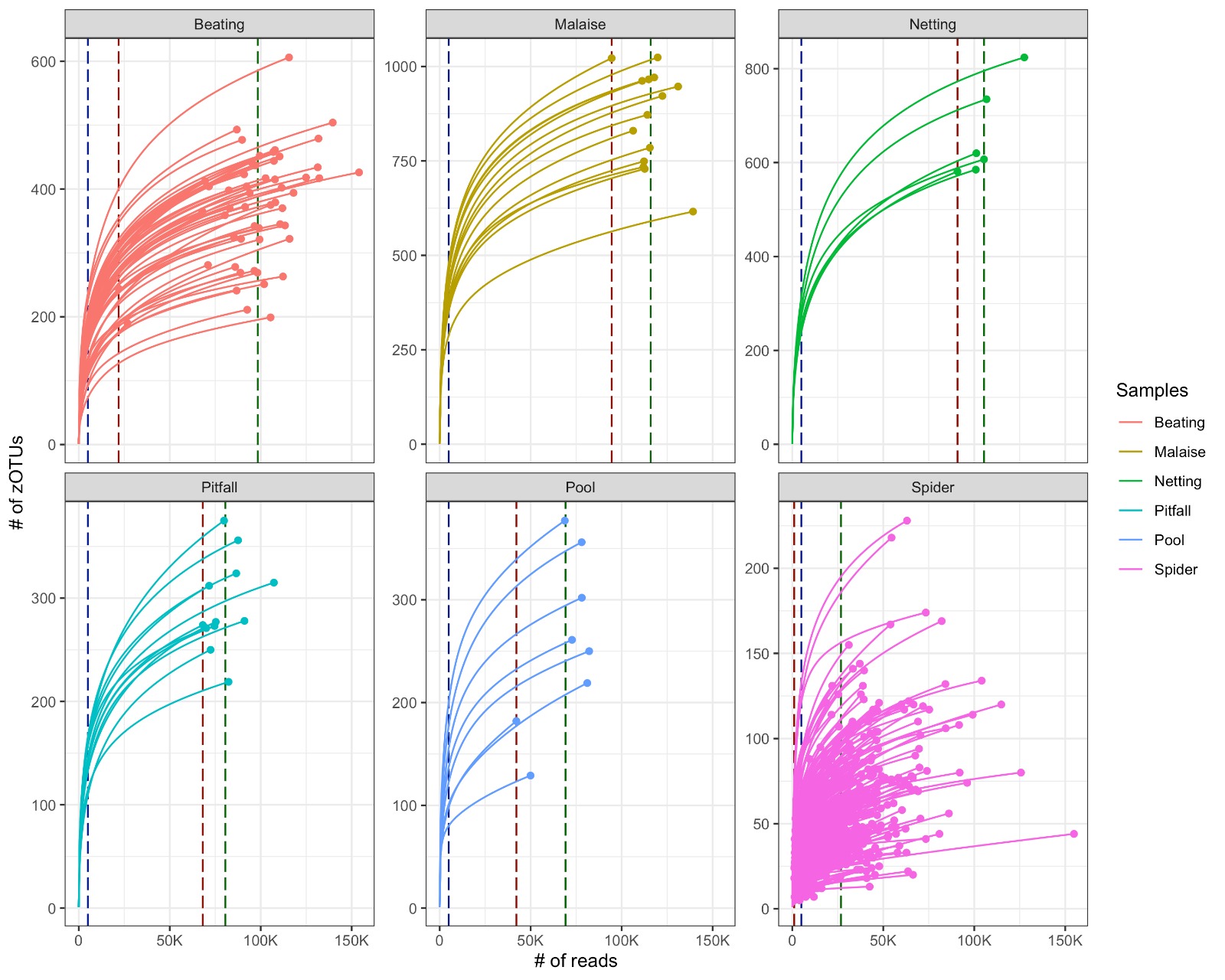

**Supplemental Figure 1: Sample saturation curves**. Each method showing each sample and the number of zOTUs are detected at each number of (top left in red) vegetation beating samples (top middle in brown) Malaise trap samples, (top right green) sweep netting samples, (bottom left teal) pitfall trap samples, (bottom middle light blue) pooled spider samples, (bottom right pink) individual spiders. Blue dashed vertical lines show 5000 read marker. Red dashed vertical lines show minimum reads per sample type. Green dashed vertical lines indicate mean reads per sample type, underlining that traditional samples were sequenced much deeper than spiders, to account for biases.

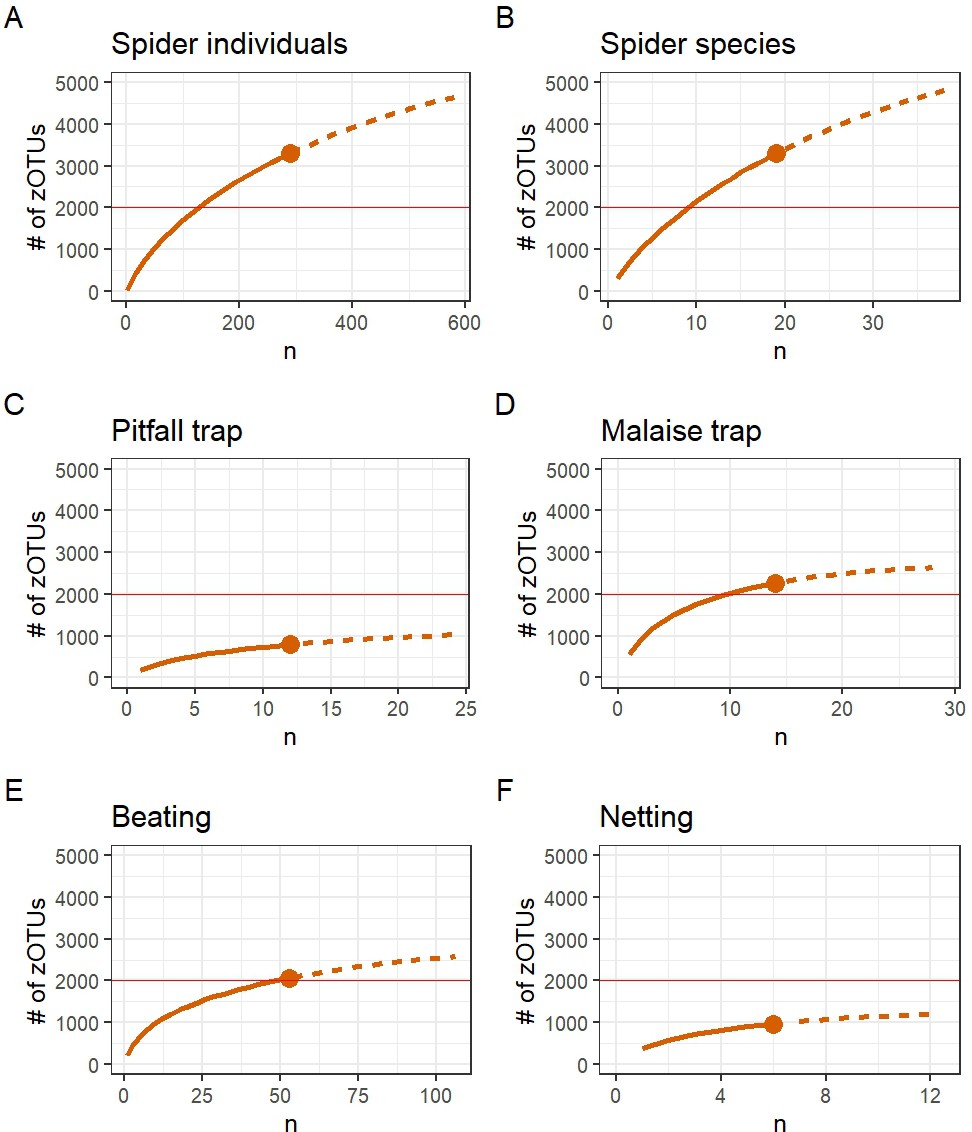

**Supplemental Figure 2: Combined Saturation curves**. Each method showing the number of zOTUs are detected at each number of (A) spider individuals, (B) spider species with n >= 10 individuals,

(C) pitfall trap samples, (D) Malaise trap samples, (E) vegetation beating samples or (F) netting samples. Dashed lines show extrapolation (calculated in R using iNEXT). Red horizontal lines visualize the level of 2000 detected zOTUs for better comparison between methods.

*Supplemental table S2: Geographic plausibility of curated archived spider samples (Argiope bruennichi) and the respective prey detected. Only zOTUs with 100% identity and 71bp fragment length scores for taxonomic annotation and more than 50 reads per occurrence are referenced here. Green color indicates plausible occurrence of prey item (source: GBIF) in the distribution area of the sample origin.*

| query | perc_i | sbjct_ | species_bold |
| --- | --- | --- | --- |
| OTU1 | 100 | 71 | Chorthippus brunneus |
| OTU1036 | 100 | 71 | Polistes dominula |
| OTU1037 | 100 | 71 | Amblyptilia acanthadactyla |
| OTU106 | 100 | 71 | Sarcophaga crassimargo |
| OTU131 | 100 | 71 | Schistocerca gregaria |
| OTU135 | 100 | 71 | Pseudochorthippus parallelus |
| OTU136 | 100 | 71 | Nysius graminicola |
| OTU14 | 100 | 71 | Apis mellifera |
| OTU148 | 100 | 71 | Pollenia rudis |
| OTU155 | 100 | 71 | Pieris rapae |
| OTU174 | 100 | 71 | Physiphora alceae |
| OTU179 | 100 | 71 | Anthomyia liturata |
| OTU190 | 100 | 71 | Philhygra palustris |
| OTU194 | 100 | 71 | Syritta pipiens |
| OTU217 | 100 | 71 | Chrysoperla nipponensis |
| OTU223 | 100 | 71 | Pherbina coryleti |
| OTU231 | 100 | 71 | Agriphila tristella |
| OTU240 | 100 | 71 | Fieberiella florii |
| OTU256 | 100 | 71 | Entomobrya multifasciata |
| OTU265 | 100 | 71 | Coenosia tigrina |
| OTU266 | 100 | 71 | Caradrina flavirena |
| OTU289 | 100 | 71 | Locusta migratoria |
| OTU305 | 100 | 71 | Toya propinqua |
| OTU311 | 100 | 71 | Asphondylia micromeriae |
| OTU340 | 100 | 71 | Scopula minorata |
| OTU36 | 100 | 71 | Tipula paludosa |
| OTU362 | 100 | 71 | Cicadella viridis |
| OTU374 | 100 | 71 | Poecilus cupreus |
| OTU422 | 100 | 71 | Helina evecta |
| OTU456 | 100 | 71 | Oulema duftschmidi |
| OTU479 | 100 | 71 | Oncocera semirubella |
| OTU48 | 100 | 71 | Pseudochorthippus parallelus |
| OTU496 | 100 | 71 | Longitarsus succineus |
| OTU5 | 100 | 71 | Andrena dorsata |
| OTU507 | 100 | 71 | Stenus impressus |
| OTU517 | 100 | 71 | Toya propinqua |
| OTU53 | 100 | 71 | Ribautodelphax collina |
| OTU558 | 100 | 71 | Aphrodes makarovi |
| OTU601 | 100 | 71 | Eriothrix rufomaculata |
| OTU7 | 100 | 71 | Pherbina coryleti |
| OTU728 | 100 | 71 | Coccinella septempunctata |
| OTU755 | 100 | 71 | Emmelina monodactyla |
| OTU76 | 100 | 71 | Sphaerophoria scripta |
| OTU78 | 100 | 71 | Tipula paludosa |
| OTU803 | 100 | 71 | Agriphila geniculea |
| OTU804 | 100 | 71 | Tephritis formosa |
| OTU913 | 100 | 71 | Gabrius nigritulus |
| OTU96 | 100 | 71 | Triodia sylvina |
| OTU97 | 100 | 71 | Luperina testacea |

| Portugal | Portugal | Spain | Spain | France | France | France | France | Belgium | Italy | Italy | Switzerland | Switzerland | Germany | Germany | Germany | Germany | Germany | Germany |
| --- | --- | --- | --- | --- | --- | --- | --- | --- | --- | --- | --- | --- | --- | --- | --- | --- | --- | --- |
| PoCeFA01P2 | PoCeGA06P1 | SpCaMB07S7 | SpCaTA14S8 | FrBnAA13 | FrCrCF27 | FrCrCP01 | FrPlAB05 | BeWANA09 | 2812 | AE2536I7 | ChGeAA04 | ChGeAA08 | GeNsBA06 | GeNwWA02 | GeNwWA05 | GeShAA04 | GeShAA06 | GeShAA08 |
| 1 | 0 | 0 | 0 | 0 | 0 | 0 | 0 | 0 | 0 | 0 | 0 | 0 | 0 | 0 | 0 | 0 | 0 | 0 |
| 0 | 0 | 0 | 0 | 0 | 0 | 0 | 0 | 0 | 0 | 1 | 0 | 0 | 0 | 0 | 0 | 0 | 0 | 0 |
| 0 | 0 | 0 | 0 | 0 | 1 | 0 | 0 | 0 | 0 | 0 | 0 | 0 | 0 | 0 | 0 | 0 | 0 | 0 |
| 0 | 0 | 0 | 0 | 0 | 0 | 0 | 0 | 0 | 0 | 0 | 0 | 0 | 0 | 0 | 0 | 1 | 0 | 0 |
| 0 | 0 | 1 | 1 | 0 | 0 | 0 | 0 | 0 | 0 | 0 | 0 | 0 | 0 | 0 | 0 | 0 | 0 | 0 |
| 0 | 0 | 0 | 0 | 0 | 0 | 0 | 0 | 1 | 0 | 0 | 0 | 0 | 0 | 0 | 0 | 0 | 0 | 0 |
| 0 | 0 | 0 | 0 | 0 | 0 | 1 | 0 | 0 | 0 | 1 | 0 | 0 | 0 | 0 | 0 | 0 | 0 | 0 |
| 0 | 0 | 0 | 0 | 0 | 1 | 1 | 0 | 0 | 0 | 0 | 0 | 0 | 0 | 1 | 0 | 0 | 0 | 0 |
| 0 | 0 | 0 | 0 | 0 | 0 | 0 | 0 | 0 | 0 | 0 | 0 | 0 | 0 | 1 | 0 | 0 | 0 | 0 |
| 0 | 0 | 0 | 0 | 0 | 0 | 0 | 0 | 0 | 1 | 1 | 0 | 0 | 0 | 0 | 0 | 0 | 0 | 0 |
| 0 | 0 | 0 | 0 | 0 | 0 | 0 | 0 | 0 | 0 | 0 | 1 | 0 | 0 | 0 | 0 | 0 | 0 | 0 |
| 0 | 0 | 0 | 0 | 0 | 1 | 0 | 0 | 0 | 0 | 0 | 0 | 0 | 0 | 0 | 0 | 0 | 0 | 0 |
| 0 | 0 | 0 | 0 | 0 | 0 | 0 | 0 | 1 | 0 | 0 | 0 | 0 | 0 | 0 | 0 | 0 | 0 | 0 |
| 0 | 0 | 0 | 0 | 0 | 0 | 0 | 0 | 0 | 0 | 0 | 0 | 0 | 0 | 1 | 0 | 0 | 0 | 0 |
| 0 | 0 | 0 | 0 | 0 | 1 | 0 | 0 | 0 | 0 | 0 | 0 | 0 | 0 | 0 | 0 | 0 | 0 | 0 |
| 0 | 0 | 0 | 0 | 0 | 0 | 0 | 0 | 0 | 0 | 0 | 0 | 0 | 0 | 0 | 0 | 1 | 0 | 0 |
| 0 | 0 | 0 | 0 | 0 | 0 | 0 | 1 | 0 | 0 | 0 | 0 | 0 | 0 | 0 | 0 | 0 | 0 | 0 |
| 0 | 1 | 0 | 0 | 0 | 0 | 0 | 0 | 0 | 0 | 0 | 0 | 0 | 0 | 0 | 0 | 0 | 0 | 0 |
| 0 | 0 | 0 | 0 | 0 | 0 | 0 | 0 | 0 | 0 | 0 | 0 | 0 | 1 | 0 | 0 | 0 | 0 | 0 |
| 0 | 0 | 0 | 0 | 0 | 0 | 0 | 0 | 0 | 0 | 0 | 0 | 0 | 0 | 0 | 0 | 1 | 0 | 0 |
| 0 | 0 | 0 | 0 | 0 | 0 | 0 | 1 | 0 | 0 | 0 | 0 | 0 | 0 | 0 | 0 | 0 | 0 | 0 |
| 0 | 0 | 1 | 0 | 0 | 0 | 0 | 0 | 0 | 0 | 0 | 0 | 0 | 0 | 0 | 0 | 0 | 0 | 0 |
| 0 | 0 | 0 | 0 | 0 | 1 | 0 | 0 | 0 | 0 | 0 | 0 | 0 | 0 | 0 | 0 | 0 | 0 | 0 |
| 0 | 0 | 0 | 0 | 0 | 1 | 0 | 0 | 0 | 0 | 0 | 0 | 0 | 0 | 0 | 0 | 0 | 0 | 0 |
| 0 | 0 | 0 | 0 | 0 | 0 | 0 | 0 | 0 | 1 | 1 | 0 | 0 | 0 | 0 | 0 | 0 | 0 | 0 |
| 0 | 0 | 0 | 0 | 0 | 0 | 0 | 0 | 0 | 0 | 0 | 0 | 0 | 0 | 0 | 0 | 1 | 1 | 1 |
| 1 | 0 | 0 | 0 | 0 | 0 | 0 | 0 | 0 | 0 | 0 | 0 | 0 | 0 | 0 | 0 | 0 | 0 | 0 |
| 0 | 0 | 0 | 0 | 0 | 0 | 0 | 0 | 1 | 0 | 0 | 0 | 0 | 0 | 0 | 0 | 0 | 0 | 0 |
| 1 | 0 | 0 | 0 | 0 | 0 | 0 | 0 | 0 | 0 | 0 | 0 | 0 | 0 | 0 | 0 | 0 | 0 | 0 |
| 0 | 0 | 0 | 0 | 0 | 0 | 0 | 0 | 0 | 0 | 0 | 0 | 0 | 1 | 0 | 0 | 0 | 0 | 0 |
| 0 | 0 | 0 | 0 | 0 | 0 | 0 | 0 | 0 | 0 | 0 | 0 | 1 | 0 | 0 | 0 | 0 | 0 | 0 |
| 0 | 0 | 0 | 0 | 0 | 0 | 0 | 0 | 1 | 0 | 0 | 0 | 0 | 1 | 0 | 0 | 0 | 0 | 0 |
| 0 | 0 | 0 | 0 | 0 | 0 | 0 | 0 | 0 | 0 | 0 | 0 | 0 | 0 | 0 | 1 | 0 | 0 | 0 |
| 0 | 0 | 0 | 0 | 0 | 0 | 0 | 0 | 0 | 0 | 0 | 0 | 0 | 0 | 1 | 0 | 0 | 0 | 0 |
| 0 | 0 | 0 | 0 | 1 | 0 | 0 | 0 | 0 | 0 | 0 | 0 | 0 | 0 | 0 | 0 | 0 | 0 | 0 |
| 0 | 0 | 0 | 0 | 0 | 0 | 0 | 0 | 0 | 0 | 1 | 0 | 0 | 0 | 0 | 0 | 0 | 0 | 0 |
| 0 | 0 | 0 | 0 | 0 | 0 | 0 | 0 | 0 | 0 | 0 | 0 | 0 | 0 | 0 | 0 | 0 | 0 | 1 |
| 0 | 0 | 0 | 0 | 0 | 0 | 0 | 0 | 0 | 0 | 0 | 0 | 0 | 0 | 0 | 1 | 0 | 0 | 0 |
| 0 | 0 | 0 | 0 | 0 | 0 | 0 | 0 | 1 | 0 | 0 | 0 | 0 | 0 | 0 | 0 | 0 | 0 | 0 |
| 0 | 0 | 0 | 0 | 0 | 0 | 0 | 0 | 0 | 0 | 0 | 0 | 0 | 0 | 0 | 0 | 1 | 0 | 0 |
| 0 | 0 | 0 | 0 | 0 | 0 | 0 | 1 | 0 | 0 | 0 | 0 | 0 | 0 | 0 | 0 | 0 | 0 | 0 |
| 0 | 0 | 0 | 0 | 0 | 0 | 1 | 0 | 0 | 0 | 0 | 0 | 0 | 0 | 0 | 0 | 0 | 0 | 0 |
| 0 | 0 | 0 | 0 | 1 | 0 | 0 | 0 | 0 | 0 | 0 | 0 | 0 | 0 | 1 | 0 | 0 | 0 | 0 |
| 0 | 0 | 0 | 0 | 0 | 0 | 0 | 0 | 0 | 0 | 0 | 0 | 0 | 0 | 0 | 0 | 1 | 0 | 0 |
| 0 | 0 | 0 | 0 | 0 | 0 | 1 | 0 | 0 | 0 | 0 | 0 | 0 | 0 | 0 | 0 | 0 | 0 | 0 |
| 0 | 0 | 0 | 0 | 0 | 0 | 1 | 0 | 0 | 0 | 0 | 0 | 0 | 0 | 0 | 0 | 0 | 0 | 0 |
| 0 | 0 | 0 | 0 | 0 | 1 | 0 | 0 | 0 | 0 | 0 | 0 | 0 | 0 | 0 | 0 | 0 | 0 | 1 |
| 0 | 0 | 0 | 0 | 0 | 0 | 0 | 0 | 0 | 0 | 0 | 0 | 0 | 0 | 0 | 0 | 1 | 0 | 0 |
| 0 | 1 | 0 | 0 | 0 | 0 | 0 | 0 | 0 | 0 | 0 | 0 | 0 | 0 | 0 | 0 | 0 | 0 | 0 |

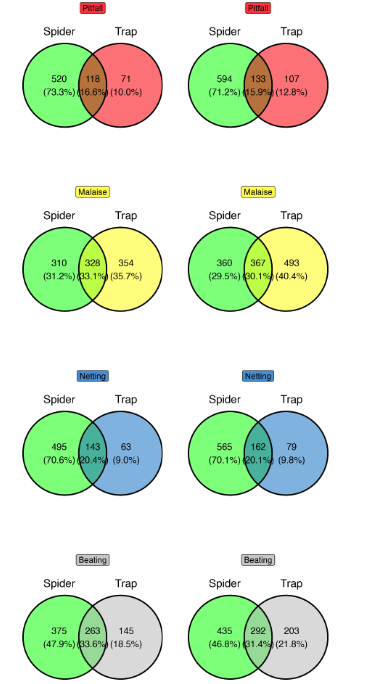

**Supplemental Figure 3. Ground-truthing Experiment 1: spider guts vs. traps at the genus level.** Venn diagrams show the overlap of detected genera between spider gut-content samples (“Spider”) and paired trap samples (“Trap”) for each method (pitfall, Malaise, netting, beating). Left column: zOTU clustering; right column: 97% OTU clustering. Numbers are counts of genera (percentages within circles), with the intersection indicating shared genera. Overall, genus-level overlap between spiders and traps is substantial across methods, and the qualitative pattern is consistent between zOTU and OTU clustering, with only minor shifts in the proportions unique to each source.

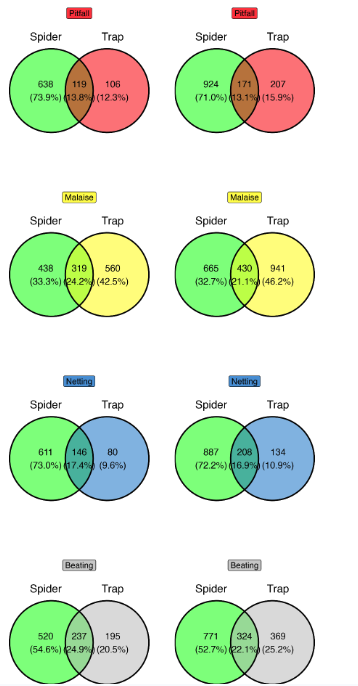

**Supplemental Figure 4. Ground-truthing Experiment 1 after collapsing ASVS/zOTUs and OTUs to taxon proxies via BLAST**. Venn diagrams compare detections from spider gut-content samples (“Spider”) and paired trap samples (“Trap”) after mapping each zOTU/OTU to a single taxon proxy using BLAST: species names assigned at ≥98% identity (with sufficient alignment coverage); otherwise the record was mapped to the most specific available rank (genus, then family, then order), labeled as “<rank> spec.” (e.g., Argiope spec., Lycosidae spec., Araneae spec.). Panels show results at the species-proxy (left column) and genus-proxy (right column) levels; companion plots provide family and order. Numbers are counts of proxies; percentages within circles indicate the fraction unique to each source, with the intersection denoting shared proxies. Collapsing reduces inflation from micro-variants/OTUs while preserving a shared core of prey taxa between spiders and traps and revealing method-specific complements.

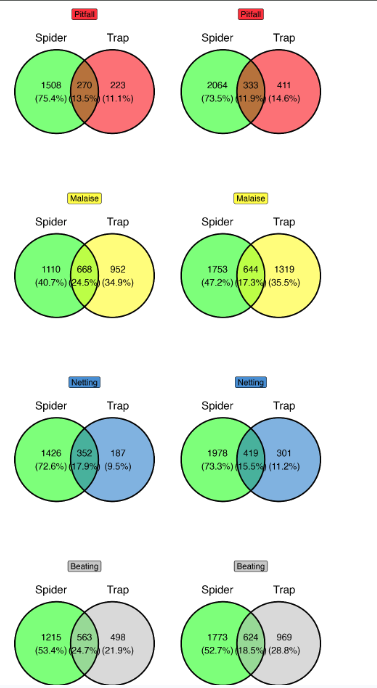

**Supplemental Figure 5. Ground-truthing Experiment 1 at the sequence-variant level (ASV/zOTU vs. 97% OTU).** Venn diagrams compare detections from spider gut-content samples (“Spider”) and paired trap samples (“Trap”) for each method (pitfall, Malaise, netting, beating). **Left column:** ASV (zOTU) level. **Right column:** 97% OTU level. Numbers are counts of detected units; percentages within circles refer to the fraction unique to each source, and the intersection denotes shared detections. Overall, overlap between spiders and traps is substantial across methods, while absolute counts shift with the clustering scheme; the qualitative pattern (a shared core plus method-specific complements) is consistent in both ASV and OTU views.

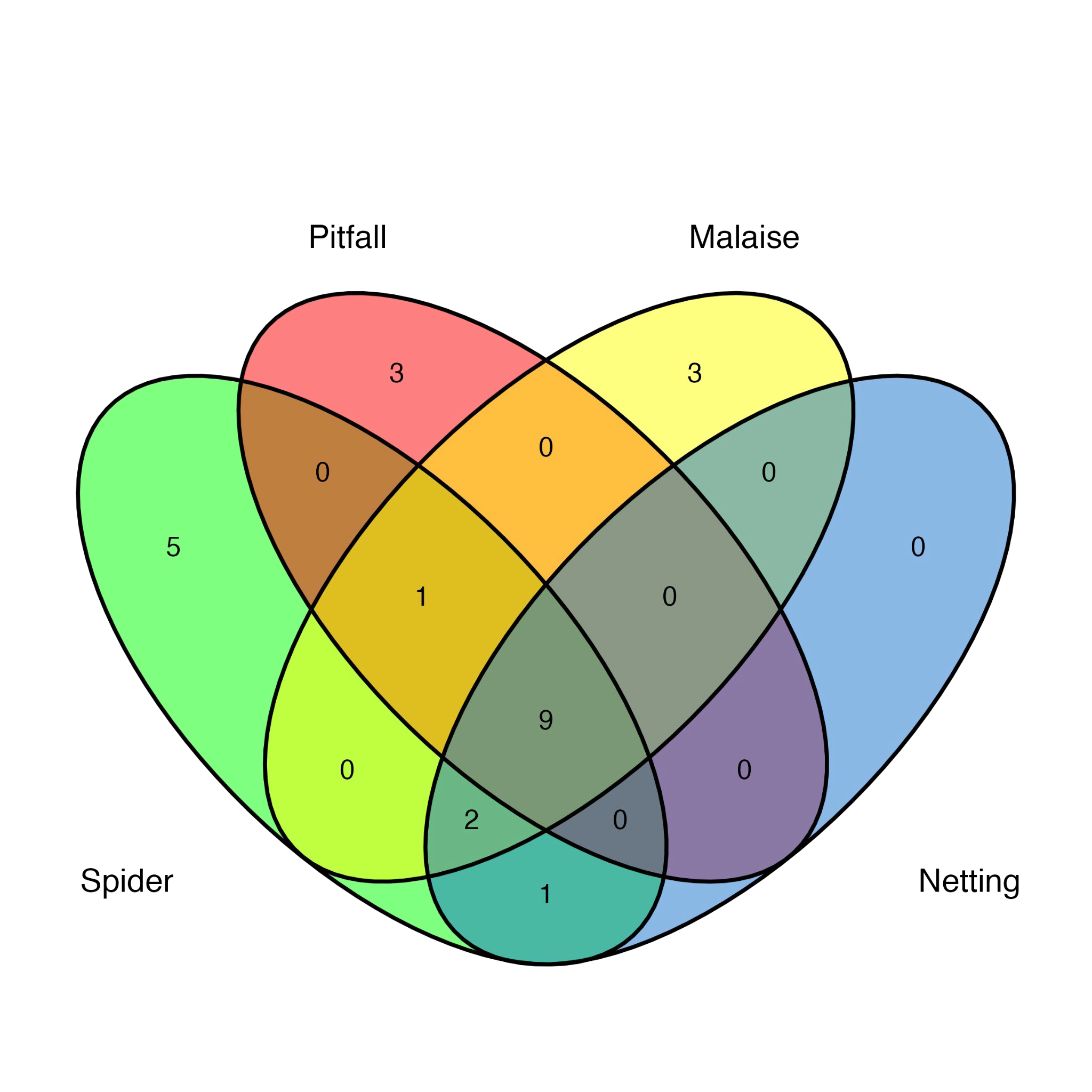

**Supplemental Figure 6: Venn diagram**. Based on the presence/absence on order level comparing the number of orders unique to and shared between each tested traditional method (red = Pitfall trap, n = 45; yellow = Malaise trap, n = 12; blue = sweep netting, n = 6) against the spider gut content (green, n = 290)

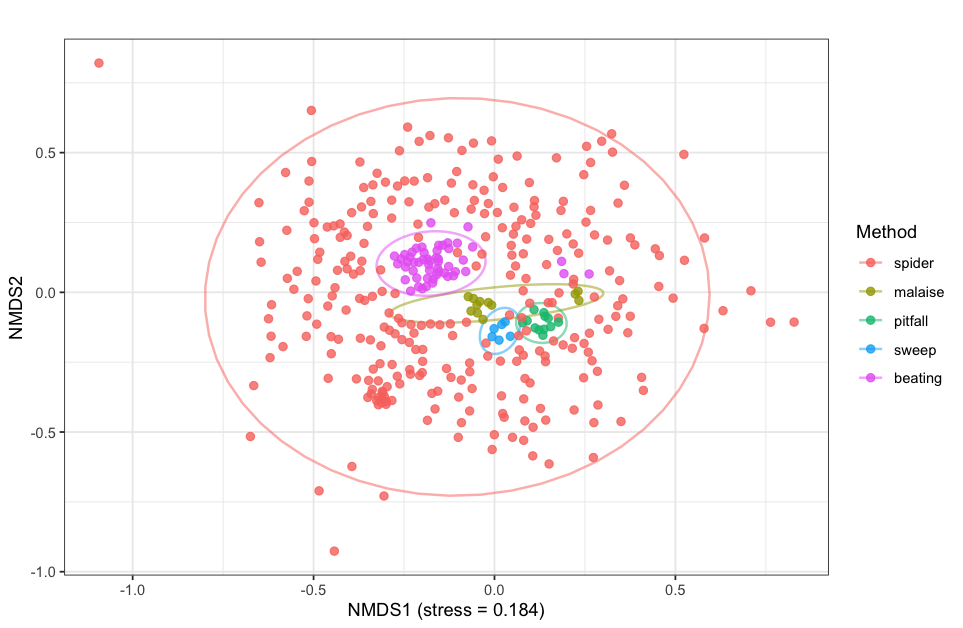

**Supplemental Figure 7** NMDS ordination of arthropod community composition recovered by spider gut content analysis and four conventional sampling methods (Malaise trap, pitfall trap, sweep netting, vegetation beating) in Experiment 1A (Germany), based on Bray–Curtis dissimilarities (stress = 0.184). Each point represents one sample. Spider gut content samples occupy a broader area of ordination space and overlap with all traditional sampling methods, whereas individual trapping methods form more compact, method-specific clusters. Ellipses indicate the dispersion of samples within each method.

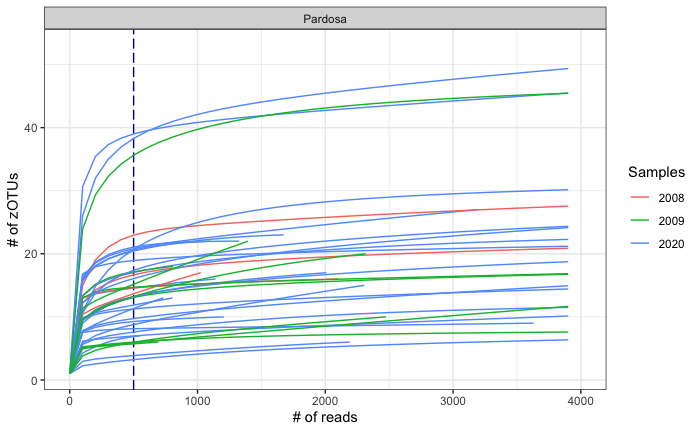

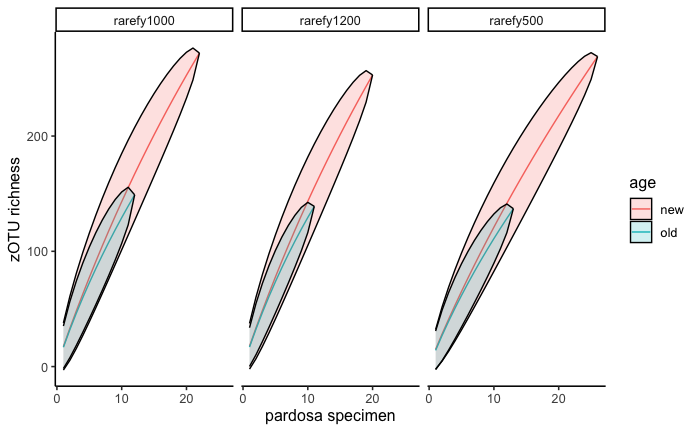

**Supplemental Figure 8. Sequencing depth and richness recovery in Pardosa specimens.** (TOP) Rarefaction curves showing zOTU accumulation as a function of sequencing depth across three sampling years. Lines represent individual specimens, colored by year. The dashed line indicates the standardized read depth used for downstream rarefaction.
(BOTTOM) Comparison of zOTU richness between newly collected and older Pardosa specimens under three rarefaction thresholds (500, 1000, 1200 reads). Ellipses illustrate the distribution of richness among age classes, showing consistently lower richness in older samples regardless of depth.
